# Cytoplasmic mRNA levels are regulated by a combination of chromatin retention and RNA stability

**DOI:** 10.1101/2022.10.24.513557

**Authors:** Callum Henfrey, Shona Murphy, Michael Tellier

## Abstract

Transcription and co-transcriptional processes, including pre-mRNA splicing and mRNA cleavage and polyadenylation, regulate the production of mature mRNAs. The carboxyl terminal domain (CTD) of RNA polymerase (pol) II, which comprises 52 repeats of the Tyr1Ser2Pro3Thr4Ser5Pro6Ser7 peptide, is involved in the coordination of transcription with co-transcriptional processes. The pol II CTD is dynamically modified by protein phosphorylation, which regulates recruitment of transcription and co-transcriptional factors. We have investigated whether cytoplasmic levels of mature mRNA from intron-containing protein-coding genes are related to pol II CTD phosphorylation, RNA stability, and pre-mRNA splicing and mRNA cleavage and polyadenylation efficiency. We find that genes that produce a low level of mature mRNA are associated with relatively high phosphorylation of the pol II CTD Tyr1 and Thr4 residues, poor RNA processing, increased chromatin retention, and shorter RNA half-life. While these poorly-processed transcripts are degraded by the nuclear RNA exosome, our results indicate that in addition to RNA half-life, chromatin retention due to a low RNA processing efficiency also plays an important role in the regulation of cytoplasmic mRNA levels.

## INTRODUCTION

Transcription of human protein coding genes by RNA polymerase (pol) II is a highly complex process requiring the coordination of multiple proteins. In addition to the transcription cycle, composed of transcription initiation, pol II pausing and release, transcription elongation, and transcription termination, co-transcriptional processes, including capping, splicing, cleavage and polyadenylation, and co-transcriptional loading of mRNA export factors are required for the production of mature mRNA (1,2). A key element regulating the crosstalk between transcription and co-transcriptional processes is the carboxyl terminal domain (CTD) of the large subunit of pol II, which comprises 52 repeats of the heptapeptide Tyr1-Ser2-Pro3-Thr4-Ser5-Pro6-Ser7. The pol II CTD can be modified by several post-translational modifications (PTMs), including protein phosphorylation, methylation, acetylation, and proline isomerization (3,4). Phosphorylation of the pol II CTD is one of the major PTMs and can occur on five residues, Tyr1, Ser2, Thr4, Ser5, and Ser7 (3,4). Kinases, including several cyclin-dependent kinases (CDKs), and phosphatases regulate the CTD phosphorylation pattern and level across the transcription cycle (3,4).

Phospho-Ser5 (Ser5P) and phospho-Ser2 (Ser2P) are the most studied modifications and are found at the promoter region and in the gene body/downstream of the poly(A) site, respectively (4). Ser5P is associated with the recruitment of the mRNA capping complex and pre-mRNA splicing factors while Ser2P is linked to the recruitment of elongation factors and proteins of the mRNA cleavage and polyadenylation complex (CPA). The roles of the three other residues, Tyr1, Thr4, and Ser7, are less well understood (5). On protein-coding genes, phosphorylation of Tyr1 is present at promoter and transcription termination regions. Tyr1P has been found to be higher on antisense promoters (PROMPTs) and enhancers compared to protein-coding genes (6,7) and to increase following DNA double-strand breaks and UV irradiation (8,9). In addition, mutation of three quarters of the tyrosine residues to alanine promotes transcriptional readthrough and a loss of the Mediator and Integrator complexes from pol II. Phosphorylation of Thr4 is found at the 3’end of protein-coding genes, indicating a potential role in transcription termination (10,11). In addition, Thr4P has been found to be higher on the gene bodies of long non-coding (lnc)RNAs, which are known to be prone to premature transcription termination (PTT) (10,11). Mutation of Thr4 residues to alanine results in a transcription elongation defect on protein-coding genes and a 3’end-processing defect of histone gene transcripts (10,12). Phosphorylation of Ser7 is currently the least understood but follows a similar pattern to Ser2P. The combination of Ser2P and Ser7P has been shown to be involved in the recruitment of the Integrator complex to small nuclear (sn)RNA genes (13). Mutation of Ser7 residues to alanine results in a decreased transcription of snRNA genes and 3’ processing of transcripts while protein coding genes do not seem to be affected (13,14).

Modification of the pol II CTD is therefore critical for coordinating co-transcriptional processes during transcription. In turn, co-transcriptional processes can affect transcription (2). A major example is the coupling between pre-mRNA splicing, mRNA cleavage and polyadenylation, and transcription termination. It has been shown via long-read sequencing approaches that protein-coding genes transcripts that are poorly processed are associated with pol II transcriptional readthrough, likely due to a failure to recognize the poly(A) site (15,16). Additionally, transcriptional readthrough can be promoted by knockdown of CPA factors, cellular stresses, or viral infection (17-19).

While important steps in the regulation of expression of protein-coding genes occur at the 5’ end of genes, including transcription initiation and pol II pause release, it is becoming increasingly clear that premature transcription termination (PTT) is a major regulator of gene expression. PTT can happen across the whole gene unit, from pol II pausing sites to poly(A) sites, mediated by mRNA decapping followed by Xrn2 degradation (20), the Integrator complex (21-24), co-transcriptional recruitment of the RNA exosome (25), U1 telescripting and intronic poly(A) site usage (26-30), and at the poly(A)-associated checkpoint (31-34).

To better understand how transcription and co-transcriptional processes regulate the production of mature mRNAs from intron-containing protein-coding genes (termed protein-coding genes in the rest of the manuscript), we took advantage of genome-wide data available for HeLa cells. We find that pol II on protein-coding genes producing relatively low levels of mature mRNA is hyperphosphorylated on the CTD Thr4, and to a lesser extent Tyr1, residues. Interestingly, the reduced production of mature mRNAs from these protein-coding genes is mediated by poor RNA processing, which results in a combination of chromatin retention of the transcripts and degradation by the nuclear RNA exosome of the poorly-processed transcripts that are released from chromatin.

Our results indicate that the regulation of RNA processing efficiency plays an important role in controlling gene expression through chromatin retention of transcripts and degradation by the nuclear RNA exosome of chromatin-released transcripts. This RNA processing efficiency-dependent regulation of mRNA levels by a combination of chromatin retention and degradation by the nuclear RNA exosome is shared between lncRNAs and protein-coding genes, indicating a general regulatory mechanism.

## MATERIAL AND METHODS

### Genome-wide datasets

The genome-wide data used in this study are summarized in Table 1.

**Table 1.** List of genome-wide data and their associated GEO accession numbers and references used in this study.

| Sample name (number of biological replicates) | GEO study | Reference |
| --- | --- | --- |
| HeLa Chromatin RNA-seq (5) | GSE81662, GSE110028 | (11,54) |
| HeLa Nucleoplasm RNA-seq (4) | GSE81662, GSE110028 | (11,54) |
| HeLa Cytoplasmic RNA-seq (2) | GSE110028 | (54) |
| HeLa Chromatin RNA-seq siEXOSC3 and associated siLuc (2) | GSE81662 | (11) |
| HeLa Nucleoplasm RNA-seq siEXOSC3 and siLuc (2) | GSE81662 | (11) |
| HeLa Chromatin RNA-seq siCPSF73 and associated siLuc (2) | GSE60358 | (40) |
| HeLa Total pol II mNET-seq siEXOSC3 and associated siLuc (2) | GSE81662 | (11) |
| HeLa Total pol II mNET-seq (2) | GSE60358, GSE81662 | (11,40) |
| HeLa Pol II Y1P mNET-seq, Empigen treated (2) | GSE81662 | (11) |
| HeLa Pol II S2P mNET-seq,<br>Empigen treated (2) | GSE81662 | (11) |
| HeLa Pol II T4P mNET-seq,<br>Empigen treated (2) | GSE81662 | (11) |
| HeLa Pol II S5P mNET-seq,<br>Empigen treated (2) | GSE81662 | (11) |
| HeLa Pol II S7P mNET-seq (2) | GSE81662 | (11) |
| HeLa Total pol II POINT-seq<br>(2) | GSE159326 | (16) |
| HeLa H3K36me3 and<br>associated Input, mNuc-seq (2) | GSE110028 | (54) |
| HeLa CPSF73 and associated<br>Input, ChIP-seq (2) | GSE127256 | (29) |
| HeLa PCF11 and associated<br>Input, ChIP-seq (2) | GSE127256 | (29) |
| HeLa Xrn2 and associated<br>Input, ChIP-seq (1) | GSE36185 | (20) |
| Raji total pol II, Ser2P, Ser5P,<br>and Thr4P (1) | GSE37519 | (10) |
| Raji total pol II, Ser2P, and<br>Tyr1P (1) | GSE52914 | (6) |
| Raji chromatin and total RNA-seq (2) | GSE94330 | (51) |

### RNA-seq and POINT-seq analysis

Chromatin, nucleoplasm, and cytoplasmic RNA-seq were analysed as previously described (35). Briefly, adapters were trimmed with Cutadapt version 1.18 (36) in paired-end mode with the following options: --minimum-length 10 -q 15,10 -j 16 -A GATCGTCGGACTGTAGAACTCTGAAC -a AGATCGGAAGAGCACACGTCTGAACTCCAGTCAC. The remaining rRNA reads were removed by mapping the trimmed reads to the rRNA genes defined in the human ribosomal DNA complete repeating unit (GenBank: U13369.1) with STAR version 2.7.3a (37) and the parameters --runThreadN 16 -- readFilesCommand gunzip -c -k --outReadsUnmapped Fastx --limitBAMsortRAM 20000000000 -- outSAMtype BAM SortedByCoordinate. The unmapped reads were mapped to the human RNU2 gene or to the GRCh38.p13 reference sequence with STAR version 2.7.3a and the parameters: --runThreadN 16 --readFilesCommand gunzip -c -k --limitBAMsortRAM 20000000000 --outSAMtype BAM SortedByCoordinate. SAMtools version 1.9 (38) was used to retain the properly paired and mapped reads (-f 3) and to create strand-specific BAM files. FPKM-normalized bigwig files were created with deepTools2 version 3.4.2 (39) bamCoverage tool with the parameters -bs 10 -p max –normalizeUsing RPKM.

### mNET-seq analysis

Adapters were trimmed with Cutadapt version 1.18 in paired-end mode with the following options: -- minimum-length 10 -q 15,10 -j 16 –A GATCGTCGGACTGTAGAACTCTGAAC –a AGATCGGAAGAGCACACGTCTGAACTCCAGTCAC. Trimmed reads were mapped to the human GRCh38.p13 reference sequence with STAR version 2.7.3a and the parameters: --runThreadN 16 -- readFilesCommand gunzip -c -k --limitBAMsortRAM 20000000000 --outSAMtype BAM SortedByCoordinate. SAMtools version 1.9 was used to retain the properly paired and mapped reads (-f 3). A custom python script (40) was used to obtain the 3’ nucleotide of the second read and the strandedness of the first read. Strand-specific bam files were generated with SAMtools. FPKM-normalized bigwig files were created with deepTools2 bamCoverage tool with the parameters -bs 1 -p max –normalizeUsing RPKM.

### ChIP-seq and mNuc-seq analysis

Adapters were trimmed with Cutadapt version 1.18 in paired-end mode with the following options: -- minimum-length 10 -q 15,10 -j 16 -A GATCGTCGGACTGTAGAACTCTGAAC -a AGATCGGAAGAGCACACGTCTGAACTCCAGTCAC. Trimmed reads were mapped to the human RNU2 gene or to the GRCh38.p13 reference sequence with STAR version 2.7.3a and the parameters: --runThreadN 16 --readFilesCommand gunzip -c -k --limitBAMsortRAM 20000000000 --outSAMtype BAM SortedByCoordinate. SAMtools version 1.9 was used to retain the properly paired and mapped reads (-f 3) and to remove PCR duplicates. Reads mapping to the DAC Exclusion List Regions (accession: ENCSR636HFF) were removed with BEDtools version 2.29.2 (41). FPKM-normalized bigwig files were created with deepTools2 bamCoverage tool with the parameters -bs 10 -p max –e – normalizeUsing RPKM.

### Proteomic analysis section

The proteome data of HeLa cells were obtained from (42). To determine whether some of the protein-coding genes could be misannotated lncRNAs, we used the global human proteome data from the Human PeptideAtlas version 2023-01 database (43). The list of genes found in the RNA-seq data was compared to the list of proteins found in the global human proteomic data to keep only protein-coding genes that are supported experimentally by proteomic data.

### Differential expression analysis

For differential expression analysis, the number of aligned reads per gene was obtained with STAR – quantMode GeneCounts option during the mapping of raw reads to the human genome or with HTSeq version 1.99.2 (44). The lists of differentially expressed genes were obtained with DESeq2 version 1.30.1 (45) and apeglm version 1.18.0 (46) keeping only the genes with a fold change < -2 or > 2 and an adjusted p-value below 0.05. The comparisons between POINT-seq, chromatin RNA-seq, and nucleoplasm RNA-seq were performed by quantifying aligned reads only on exons as the co-transcriptional pre-mRNA splicing rates, and therefore the number of reads mapped on introns, differ between the three techniques (see below).

### Human gene annotation and selection of subset of genes

Gencode V38 annotation, which is based on the hg38 version of the human genome, was used to obtain the list of all protein-coding genes. Intronless and histone genes were removed to obtain intron-containing protein-coding genes. For each gene, we kept the annotation (TSS and poly(A) site) of the highest expressed transcript isoform, which was obtained with Salmon version 1.2.1 on four HeLa chromatin RNA-seq experiments. Only transcripts that are expressed (TPM > 0) in at least three of the four biological replicates were retained. The list of similarly expressed of nucleoplasm-enriched genes, chromatin-enriched genes, and non-enriched genes was generated through iterative random-subsampling to achieve subsets of 500 genes with the most similar expression level and distribution in the chromatin RNA-seq. For the total pol II mNET-seq subsets, a comparable subsampling was performed but rather than using 500 genes, we used 10% of similarly expressed genes from each category as we could not obtain a non-significant difference with 500 genes due to the difference in nascent transcription level between chromatin-enriched and nucleoplasm-enriched genes.

### Splicing efficiency

The splicing efficiency on POINT-seq and RNA-seq was calculated by first parsing each bam file to obtain the list of spliced and unspliced reads with the awk command (awk ‘/^@/ || $6 ∼ /N/’ for spliced reads and awk ‘/^@/ || $6 !∼ /N/’ for unspliced reads). The splicing efficiency was then calculated as the number of spliced reads over total reads with BEDtools multicov –s –split. The splicing efficiency of each transcript was then normalised to the number of exons.

Significant changes in splicing events following knockdown of the nuclear RNA exosome were obtained with rMATS version 4.1.2 with the options: bam file input, paired-end mode (47).

### Transcription termination index

The transcription termination index is defined as from (40), termed readthrough index in this paper: *RTI*= log_2_(*GB* / *TES* +*c*), *c*= (*Min*(*GB* /*TES*)>0 /2).

### Correlation heatmap

The mNET-seq heatmap was computed with deepTools2 multiBamSummary tool with the following parameters: bins –bs 10000 –distanceBetweenBins 0 –p max –e. The resulting matrix was plotted with deepTools2 plotCorrelation and the following parameters: --corMethod pearson --skipZeros –colorMap RdYlBu_r --plotNumbers.

### Metaprofiles, boxplots, and violin plots

Metaprofiles, boxplots, and violin plots were generated with R version 4.0.5. Quantifications have been performed across the gene body, TSS to TES. ChIP-seq and mNuc-seq metaprofiles are shown as IP / Input signal.

### Statistical tests

The statistical tests are indicated in the figures legends and were performed with R version 4.0.5.

## RESULTS

### A subset of expressed protein-coding genes produces a low amount of mature mRNA and protein

Previous studies have shown that a subset of long intergenic non-coding (linc)RNAs, named lincRNA-like protein-coding genes, are similar to mRNAs in that they undergo RNA processing and produce stable nuclear RNA. In addition, a limited number of protein-coding gene transcripts have been found to have features common to lincRNAs, including high chromatin reads relative to transcription levels and RNA exosome sensitivity (11,48). However, the RNA exosome is only one of the protein complexes regulating RNA production. We have therefore more widely investigated the amount of mature mRNA produced from intron-containing protein-coding genes in relation to transcription and chromatin retention of transcripts.

We have used chromatin and nucleoplasm RNA-seq data available from HeLa cells to identify the protein-coding genes whose transcripts are enriched in the nucleoplasm compared to the chromatin (nucleoplasm-enriched, log2(fold change) > 1, adjusted p-value < 0.05) or that are enriched on the chromatin compared to the nucleoplasm (chromatin-enriched, log2(fold change) < -1, adjusted p-value < 0.05) (Figure 1A). As the proportion of intronic reads differ between chromatin RNA-seq and nucleoplasm RNA-seq (see below), we therefore focused our analysis on exonic reads. From the DESeq2 analysis, we initially found 4,803 nucleoplasm-enriched protein-coding genes and 6,130 chromatin-enriched protein-coding genes. We removed histone and intronless genes to keep intron-containing protein-coding genes and genes that were not found to be expressed in the nucleoplasm RNA-seq. We then kept for each remaining gene the annotation (TSS and poly(A) site) of the most expressed transcript in the nucleoplasm RNA-seq to obtain a list of 4,686 transcripts from nucleoplasm-enriched genes and 4,119 genes transcripts from chromatin-enriched genes (Figure 1B). For comparison purposes, we have also defined a set of 4,140 non-enriched set of genes (no significant difference between nucleoplasm RNA-seq and chromatin RNA-seq). To determine whether transcripts from the nucleoplasm-enriched and chromatin-enriched genes are exported to the cytoplasm, we re-analysed HeLa cytoplasmic RNA-seq data, which shows efficient cytoplasmic export of transcripts from nucleoplasm-enriched genes while mRNAs from chromatin-enriched genes have a limited presence in the cytoplasm (Figure 1C). To determine whether a lower nucleoplasm and cytoplasm RNA-seq signal also results in a lower protein production, we compared our RNA-seq results with a previously-published re-analysis of HeLa proteome datasets (42). We could only match ∼5,000 genes of the RNA-seq/proteome data but found that fewer of the chromatin-enriched genes produce protein products than nucleoplasm-enriched genes, and chromatin-enriched genes produce fewer peptides of each protein (Figure 1D). As only 306 chromatin-enriched genes were found to produce proteins in the HeLa proteome datasets, we checked whether chromatin-enriched genes could be incorrectly annotated lncRNAs. We compared the RNA-seq results with a database of 2,299 proteomic experiments from the Human PeptideAtlas (43) version 2023-01 database and found that 3,442 chromatin-enriched genes (out of 4,119 genes, 83.6%) produce peptides (Supplementary Figure 1A). In contrast, 4,583 nucleoplasm-enriched genes (out of 4,686 genes, 97.8%), and 4,088 non-enriched genes (out of 4,138 genes, 98.8%) produce proteins (Supplementary Figure 1A). While these results demonstrate that most chromatin-enriched genes are transcriptionally active and have the potential to produce proteins, some of these genes could be incorrectly annotated lncRNAs. For the subsequent analyses, we removed all the chromatin-enriched, nucleoplasm-enriched, and non-enriched genes that were not experimentally supported as protein-coding genes in the Human PeptideAtlas version 2023-01 database.

**Figure 1.**
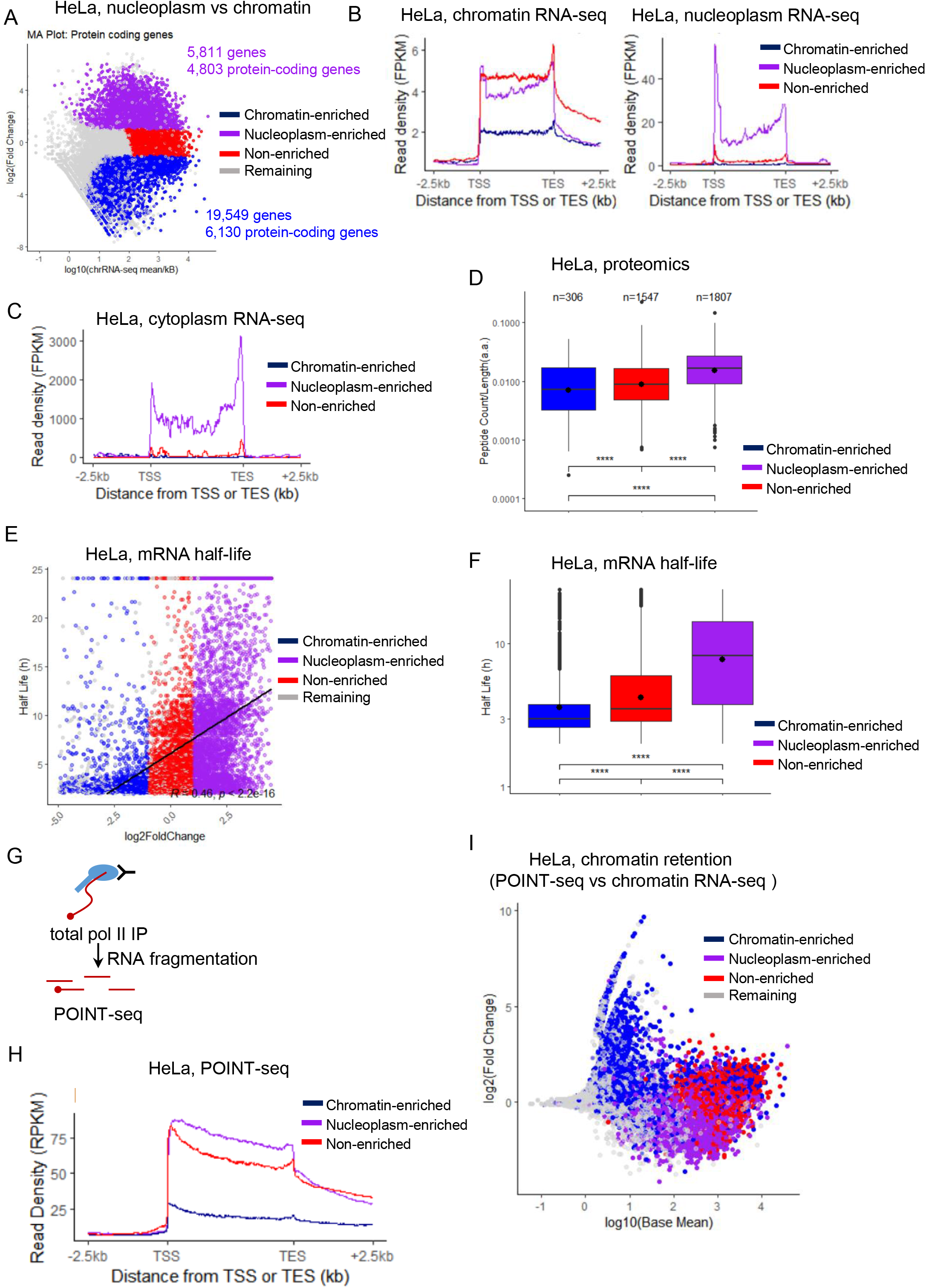
A subset of expressed protein-coding genes produces a low amount of mature mRNA and protein. **(A)** MA plot in HeLa cells of the intron-containing protein-coding genes found to be differentially enriched in the nucleoplasm (nucleoplasm-enriched, purple) or in the chromatin (chromatin-enriched, blue) fraction. The set of non-enriched genes (red) and the remaining genes (grey) are also indicated. **(B)** Chromatin RNA-seq and nucleoplasm RNA-seq metagene profiles in HeLa cells of nucleoplasm-enriched (purple), chromatin-enriched (blue), or non-enriched (red) genes. **(C)** Cytoplasmic RNA-seq metagene profiles in HeLa cells of nucleoplasm-enriched (purple), chromatin-enriched (blue), or non-enriched (red) genes. **(D)** Boxplots, shown as min to max with first quartile, median, and third quartile, of the number of peptides per protein found for nucleoplasm-enriched (purple), chromatin-enriched (blue), non-enriched (red) genes. The number of proteins found to have at least one peptide are indicated at the top of each category. Statistical test: Wilcoxon rank sum test. P-value: **** < 0.0001. **(E)** XY correlation plot of the nucleoplasm enrichment ratio (log2 fold change Nucleoplasm RNA-seq versus Chromatin RNA-seq) versus transcripts half-life obtained from (50). The Pearson correlation with p-value is indicated on the plot. Nucleoplasm-enriched (purple), chromatin-enriched (blue), non-enriched (red), and remaining (grey) genes are shown. **(F)** Boxplots, shown as min to max with first quartile, median, and third quartile, of the transcripts half-life for nucleoplasm-enriched (purple), chromatin-enriched (blue), non-enriched (red) genes. The number of proteins found to have at least one peptide are indicated at the top of each category. Statistical test: Wilcoxon rank sum test. P-value: **** < 0.0001. **(G)** Schematic of the total pol II POINT-seq experiments. **(H)** POINT-seq metagene profiles in HeLa cells of nucleoplasm-enriched (purple), chromatin-enriched (blue), or non-enriched (red) genes. **(I)** XY correlation plot of the log10 of the baseMean (average expression level) versus the log2 fold change of “Chromatin RNA-seq versus POINT-seq”, which provides a measure of chromatin retention. The Pearson correlation with p-value is indicated on the plot. Nucleoplasm-enriched (purple), chromatin-enriched (blue), non-enriched (red), and remaining (grey) genes are shown.

As chromatin-enriched genes are transcriptionally active but produce only a small amount of protein, we investigated whether the transcripts produced by the chromatin-enriched genes are more prone to degradation. We compared our RNA-seq results with a previously-published HeLa mRNA half-life dataset (49), which provides half-life data for the transcripts of 1,446 chromatin-enriched, 3,691 non-enriched, and 4,070 nucleoplasm-enriched genes (Figure 1E and F). We found a clear trend where mRNAs from chromatin-enriched genes have on average shorter half-lives compared to transcripts from non-enriched genes and nucleoplasm-enriched genes, with the latter having on average the longest half-lives.

To determine whether the chromatin enrichment of transcripts could be explained only by rapid degradation by the nuclear RNA exosome, we re-analysed HeLa POINT-seq data (16), which captures nascent RNA transcription and co-transcriptional splicing (Figure 1G and H). POINT-seq profiles of chromatin-enriched, non-enriched, and nucleoplasm-enriched genes were similar to the profiles obtained from chromatin RNA-seq (Figure 1B). Chromatin RNA-seq data represents a combination of nascent transcription and of transcripts associated with chromatin while POINT-seq data are a measure of nascent transcription, e.g. transcripts associated with pol II. We therefore reasoned that a comparison of POINT-seq and chromatin RNA-seq data over exons, which avoids technical differences on co-transcriptional splicing efficiency (see below), will indicate chromatin retention if the chromatin RNA-seq signal is enriched compared to the POINT-seq signal (Figure 1I). We found that chromatin-enriched genes are more prone to chromatin retention (blue points with a high log2 (Fold change)) compared to non-enriched (red) and nucleoplasm-enriched (purple) genes. As chromatin-enriched genes are on average longer than non-enriched and nucleoplasm-enriched genes (Supplementary Figure 1B), we investigated whether the higher chromatin retention of transcripts from chromatin-enriched genes is due to their longer size (Supplementary Figure 1C). We observed a moderate positive correlation (R = 0.32, p < 2.2e-16), indicating that gene length only partially explains the higher chromatin retention of transcripts from chromatin-enriched genes.

These findings indicate that in addition to RNA stability, the production of mature mRNAs can also be regulated by chromatin retention of transcripts.

### Higher levels of pol II Tyr1 and Thr4 phosphorylation are associated with poor expression and chromatin-enrichment of transcripts

Chromatin retention of lncRNA transcripts is associated with a different pol II CTD phosphorylation pattern and with poor co-transcriptional RNA processing, including defective pre-mRNA splicing and mRNA CPA (11). We therefore investigated whether the pol II CTD phosphorylation patterns and/or levels also differ between nucleoplasm-enriched and chromatin-enriched genes. We re-analysed HeLa mNET-seq data for total pol II and the different CTD phosphorylation marks, using Empigen-treated mNET-seq datasets when available as these identify *bone fide* mNET-seq signals without non-nascent RNA associated with the pol II (Figure 2A and Supplementary Figure 1D). While the total pol II profile follow the expected mNET-seq pattern for the three groups of genes, we observed a higher total pol II level on nucleoplasm-enriched genes compared to chromatin-enriched genes (Figure 2B and Supplementary Figure 1E and 1F). We therefore investigated whether nucleoplasm-enriched genes are more expressed (i.e. more pol II transcribing the genes) and/or pol II is slower, which will also result in a higher pol II signal. To differentiate between these possibilities, we compared the pol II elongation rate data obtained for 1,398 protein-coding genes in HeLa cells (50) to the nucleoplasm-enrichment ratio, which corresponds to log2(Fold change of Nucleoplasm RNA-seq vs Chromatin RNA-seq) (Supplementary Figure 1G). We did not observe any correlation between the pol II elongation rate and nucleoplasm-enrichment, indicating that nucleoplasm-enriched genes are not associated with slower pol II elongation and that chromatin-enriched genes are also not transcribed faster. The higher signals of total pol II, but also POINT-seq and chromatin RNA-seq (Figure 1B and 1H), on nucleoplasm-enriched genes compared to chromatin-enriched genes are therefore likely explained by a higher transcriptional level.

**Figure 2.**
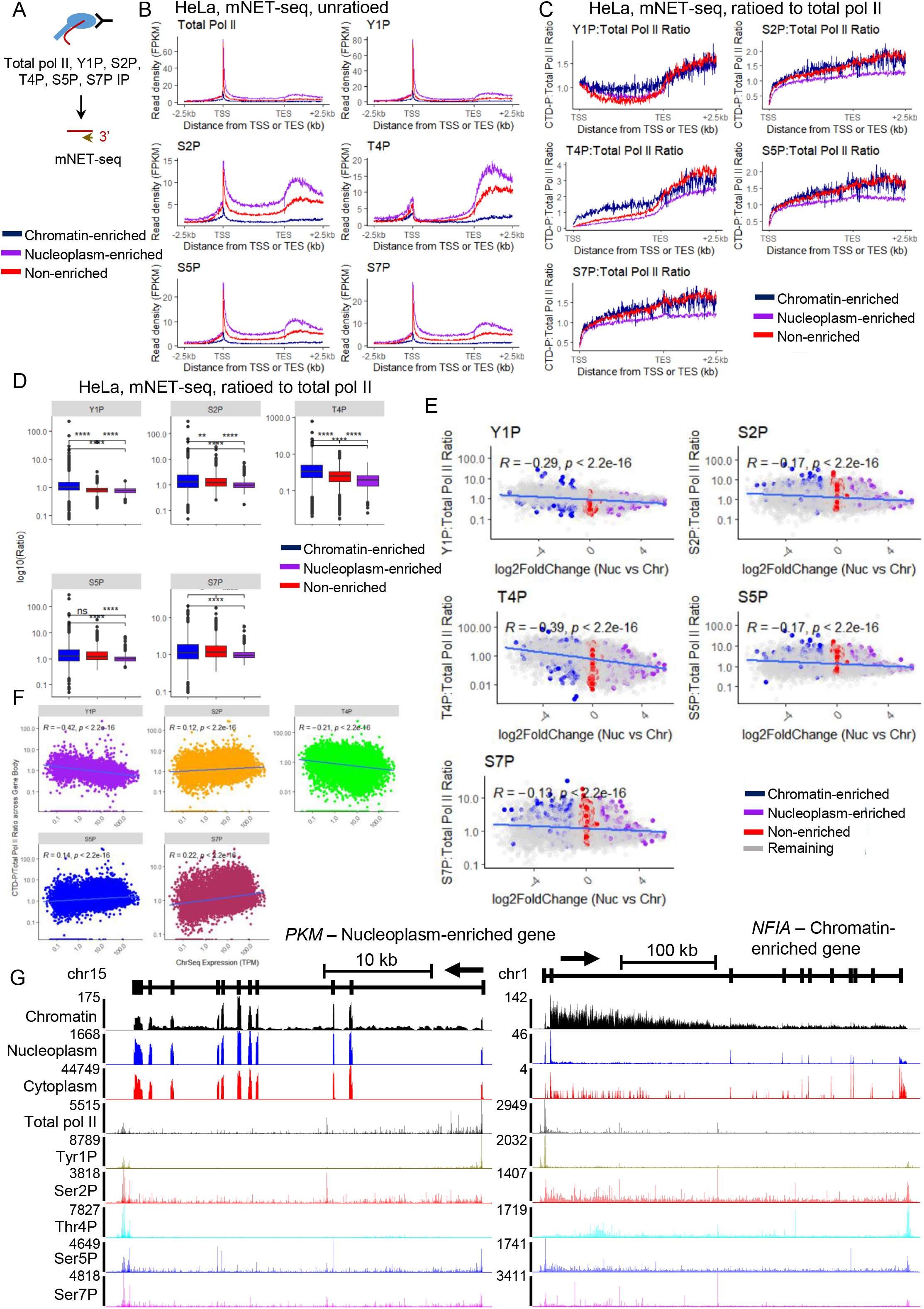
Higher levels of pol II Tyr1 and Thr4 phosphorylation are associated with poor expression and chromatin retention of transcripts. **(A)** Schematic of the total pol II mNET-seq experiments. **(B)** Metagene profiles of mNET-seq in HeLa cells of total pol II and the different pol II CTD phosphorylation mark for nucleoplasm-enriched (purple), chromatin-enriched (blue), non-enriched (red) genes. **(C)** Metagene profiles of mNET-seq in HeLa cells of each pol II CTD phosphorylation mark ratioed to total pol II for nucleoplasm-enriched (purple), chromatin-enriched (blue), non-enriched (red) genes. **(D)** Boxplots, shown as min to max with first quartile, median, and third quartile, of each pol II CTD phosphorylation mark ratioed to total pol II across the gene body of nucleoplasm-enriched (purple), chromatin-enriched (blue), non-enriched (red) genes. Statistical test: Wilcoxon rank sum test. P-value: ns: not significant, ** < 0.01, **** < 0.0001. **(E)** XY correlation plots of the nucleoplasm fold enrichment, defined as the fold change between nucleoplasm RNA-seq versus chromatin RNA-seq, and each pol II CTD phosphorylation mark ratioed to total pol II. The Pearson correlation with p-value is indicated on each plot. Nucleoplasm-enriched (purple), chromatin-enriched (blue), non-enriched (red), and remaining (grey) genes are shown. **(F)** XY correlation plots of the expression level of protein-coding gene, defined by chromatin RNA-seq, and each pol II CTD phosphorylation mark ratioed to total pol II. The Pearson correlation with p-value is indicated on each plot. **(G)** Screenshot of the genome browser chromatin RNA-seq, nucleoplasm RNA-seq, cytoplasmic RNA-seq, and total pol II and CTD phosphorylation mNET-seq tracks of the protein-coding genes *PKM* (nucleoplasm-enriched) and *NFIA* (chromatin-enriched). The arrow indicates the sense of transcription.

The different pol II CTD phosphorylation profiles follow the expected mNET-seq pattern for the three groups of genes, with a higher signal for all CTD phosphorylation marks on nucleoplasm-enriched genes compared to chromatin-enriched genes (Figure 2B) (4,11). As total pol II levels differ between the three groups of genes, we ratioed each CTD phosphorylation signal to total pol II to determine whether the CTD phosphorylation levels are similar between the three groups of genes (Figure 2C and D). We found that the nucleoplasm-enriched genes have generally less phosphorylated CTD than the chromatin-enriched genes or non-enriched genes. In contrast, Tyr1 and Thr4 phosphorylation levels are higher in the gene body of the chromatin-enriched genes compared to the non-enriched genes or the nucleoplasm-enriched genes. Interestingly, there is a negative correlation between the nucleoplasm-enrichment ratio and the gene body levels of Tyr1P (R = -0.29, p < 2.2e-16) and Thr4P (R = -0.39, p < 2.2e-16) ratioed to pol II (Figure 2E). As the total pol II level is lower on the chromatin-enriched genes compared to the nucleoplasm-enriched genes, we investigated the relationship between chromatin RNA-seq expression and gene body CTD phosphorylation ratioed to total pol II across all intron-containing protein-coding genes (Figure 2F). Although there is a weak but positive correlation between chromatin RNA-seq levels and phosphorylation of Ser2, Ser5, and Ser7, there is a negative correlation between the chromatin RNA-seq level and especially Tyr1P (R = -0.42, p < 2.2e-16) and, to a lesser extent, Thr4P (R = -0.21, p < 2.2e-16). Single gene examples of a nucleoplasm-enriched gene (*PKM*) and of a chromatin-enriched gene (*NFIA*) are shown in Figure 2G.

As Tyr1P and Thr4P levels are associated with a lower level of nascent transcription, we investigated whether the higher relative Tyr1P and Thr4P we observed for chromatin-enriched genes could be due to generally lower expression of these genes rather than a chromatin enrichment-specific CTD phosphorylation pattern (Figure 2B and Supplementary Figure 1E and F). To correct for difference in expression, we selected for each group, via an iterative random-subsampling approach, a subset of 500 genes with the most similar expression level and distribution in chromatin RNA-seq (see Methods, Figure 3A-C, Supplementary Figure 2A and B). Re-analysis of the CTD phosphorylation mNET-seq ratioed to total pol II on these three subsets of 500 genes indicates that the nucleoplasm-enriched genes have less Ser2, Thr4, Ser5, and Ser7 phosphorylation while the chromatin-enriched genes still have higher Thr4P, and to a lesser extent, Tyr1P (Figure 3D and Supplementary Figure 3C). To confirm the results, we also selected 10% of genes from each group, via an iterative random-subsampling approach, to obtain a similar nascent expression level and distribution from total pol II mNET-seq (see Methods, Figure 3E-G). Re-analysis of the CTD phosphorylation mNET-seq ratioed to total pol II on these three subsets of genes indicates that the nucleoplasm-enriched genes no longer have less Ser2, Thr4, Ser5, and Ser7 phosphorylation while the chromatin-enriched genes still have higher Tyr1 and Thr4 phosphorylation (Figure 3D-F). The subsampling performed with total pol II mNET-seq results in the selection of chromatin-enriched genes that are amongst the most expressed (compare y-axis of Figure 3E, average value around 5, to Supplementary Figure 1E, average value around 0.1). As hyperphosphorylation of the pol II CTD Thr4 residues is observed when the mark is either unratioed or ratioed to pol II for this subset of highly expressed chromatin-enriched genes, the higher Thr4 phosphorylation of chromatin-enriched genes is not simply due to generally lower expression but is a feature of this category of genes. We also investigated why the lower CTD phosphorylation to pol II ratio observed on nucleoplasm-enriched genes disappeared with the total pol II mNET-seq subsampling. Unlike the full gene set and chromatin RNA-seq subset of nucleoplasm-enriched genes, the total pol II mNET-seq subset of genes is on average not shorter than the chromatin-enriched and the non-enriched genes (Supplementary Figure 1B and Supplementary Figure 3A and 3B). However, we could not find any correlation between CTD phosphorylation ratioed to pol II and gene length (Supplementary Figure 3C), indicating another reason behind the disappearance. Comparison of the nucleoplasm-enrichment ratio with relative CTD phosphorylation reveals a correlation where the most nucleoplasm enriched genes have lower phosphorylation levels for Ser2, Ser5 and Ser5 ratioed to total pol II signal (see distribution of data points for the genes with a nucleoplasm-enrichment ratio above 4 in Figure 2E). We therefore plotted the distribution of the nucleoplasm-enrichment ratios for the different subsets (Supplementary Figure 3D). The total pol II mNET-seq subset of nucleoplasm-enriched genes shows a specific decrease in the average nucleoplasm-enrichment ratio compared to the total genes and the chromatin RNA-seq subset genes, likely explaining the disappearance of the lower total pol II ratioed CTD phosphorylation levels.

**Figure 3.**
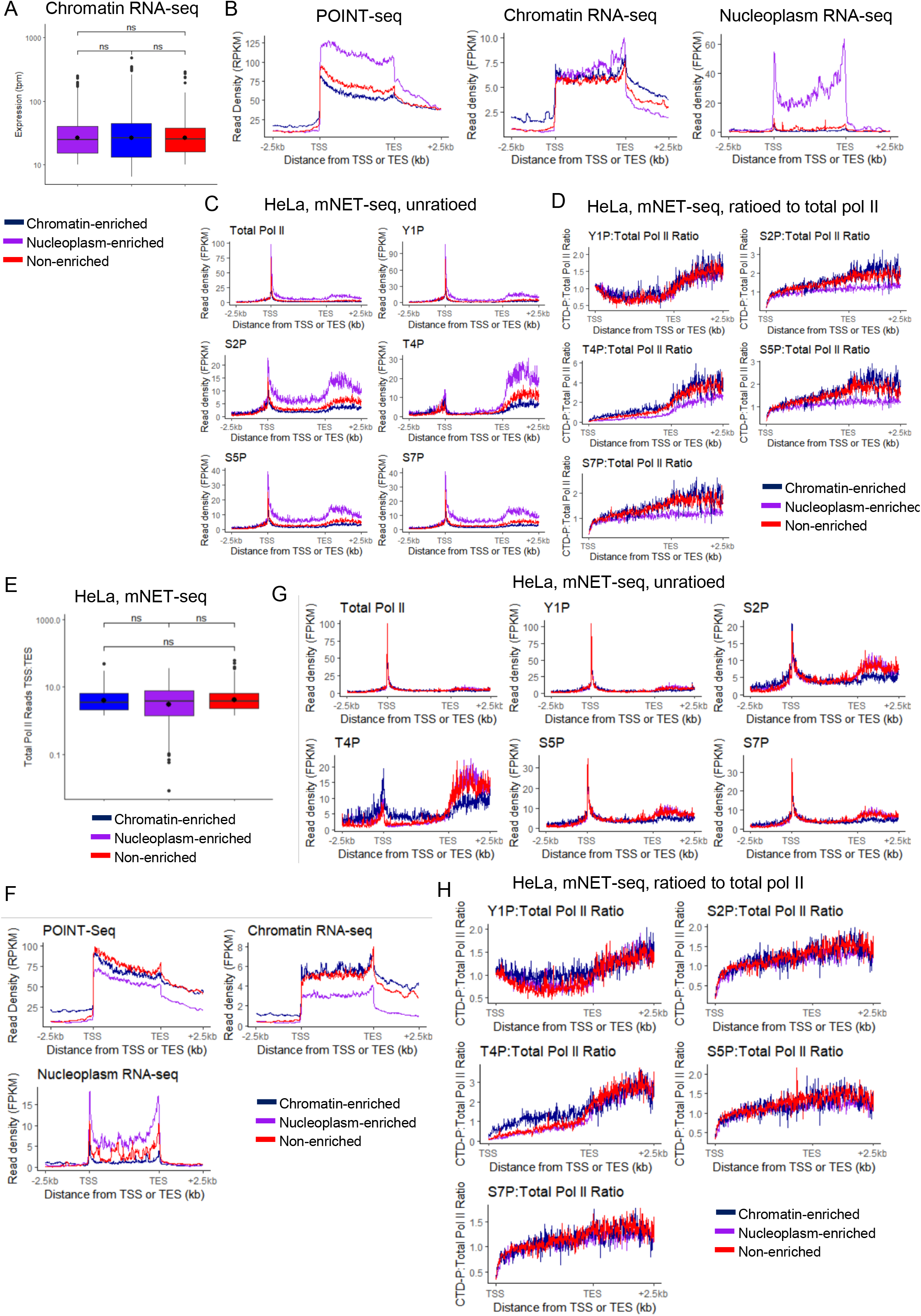
Transcripts from chromatin-enriched genes are associated with a higher phosphorylation of the pol II CTD Thr4 residues. **(A)** Boxplots, shown as min to max with first quartile, median, and third quartile, of the expression (TPM) in chromatin RNA-seq of the chromatin RNA-seq selected 500 nucleoplasm-enriched (purple), chromatin-enriched (blue), or non-enriched (red) genes. **(B)** Metagene profiles in HeLa cells of POINT-seq, chromatin RNA-seq, and nucleoplasm RNA-seq of the chromatin RNA-seq selected 500 nucleoplasm-enriched (purple), chromatin-enriched (blue), or non-enriched (red) genes. **(C)** Metagene profiles of mNET-seq in HeLa cells of total pol II and the different pol II CTD phosphorylation mark across the chromatin RNA-seq selected 500 nucleoplasm-enriched (purple), chromatin-enriched (blue), or non-enriched (red) genes. **(D)** Metagene profiles of mNET-seq in HeLa cells of each pol II CTD phosphorylation mark ratioed to total pol II across the chromatin RNA-seq selected 500 nucleoplasm-enriched (purple), chromatin-enriched (blue), or non-enriched (red) genes. **(E)** Boxplots, shown as min to max with first quartile, median, and third quartile, of the total pol II mNET-seq expression of the total pol II mNET-seq selected nucleoplasm-enriched (purple), chromatin-enriched (blue), or non-enriched (red) genes. **(F)** Metagene profiles in HeLa cells of POINT-seq, chromatin RNA-seq, and nucleoplasm RNA-seq of the total pol II mNET-seq selected nucleoplasm-enriched (purple), chromatin-enriched (blue), or non-enriched (red) genes. **(G)** Metagene profiles of mNET-seq in HeLa cells of total pol II and the different pol II CTD phosphorylation mark across the total pol II mNET-seq selected nucleoplasm-enriched (purple), chromatin-enriched (blue), or non-enriched (red) genes. **(H)** Metagene profiles of mNET-seq in HeLa cells of each pol II CTD phosphorylation mark ratioed to total pol II across the total pol II mNET-seq selected nucleoplasm-enriched (purple), chromatin-enriched (blue), or non-enriched (red) genes.

To confirm the observations made in HeLa cells, we also reanalysed chromatin RNA-seq and total RNA-seq from Raji cells (51) (Supplementary Figure 4A and B). Re-analysis of Raji pol II CTD datasets (6,10) on the groups of Raji total-enriched, non-enriched, and chromatin-enriched also indicates that there is higher Tyr1 and Thr4 phosphorylation on chromatin-enriched genes while total-enriched genes have less Tyr1, Ser2, and Thr4 phosphorylation (Supplementary Figure 4C and D). Comparison of the genes expressed in both HeLa and Raji shows that 47-63% of the nucleoplasm/total-enriched genes and 19-44% of chromatin-enriched genes are common between both cell lines (Supplementary Figure 4E). In addition, ∼160 genes were found to be in opposite categories between the two cell lines (chromatin-enriched to nucleoplasm/total-enriched or vice-versa).

These findings indicate that higher phosphorylation of Thr4, and to a lesser extent Tyr1, are markers of chromatin-enriched genes while the nucleoplasm-enriched genes with the highest nucleoplasm enrichment ratio are rather associated with lower CTD phosphorylation levels.

### Transcripts from chromatin-enriched genes are poorly processed

As pol II CTD phosphorylation is associated with co-transcriptional processes and we observed differences in CTD phosphorylation levels between nucleoplasm-enriched and chromatin-enriched genes, we investigated whether RNA processing efficiency, including pre-mRNA splicing and mRNA CPA, also differs between the two groups of genes. For pre-mRNA processing, we re-analysed HeLa POINT-seq data, which captures nascent RNA transcription and co-transcriptional splicing (Figure 1G) (16). We calculated co-transcriptional splicing efficiency as the ratio of spliced reads over total reads across each intron-containing protein-coding transcript. As expected, we observed a correlation between the number of exons and our measure of splicing efficiency (Supplementary Figure 5A), which agrees with previous observations that gene length positively correlates with co-transcriptional splicing efficiency (52). As the distribution of the number of exons per gene differs between the chromatin-enriched, nucleoplasm-enriched, and non-enriched genes (Supplementary Figure 5B), we normalized the splicing efficiency of each transcript to its number of exons (Figure 4A and B). We find for the three datasets (POINT-seq, chromatin RNA-seq, and nucleoplasm RNA-seq) that the chromatin-enriched gene transcripts have the lowest splicing efficiency while the nucleoplasm-enriched gene transcripts have the highest splicing efficiency. Importantly, while chromatin-enriched genes are longer on average than non-enriched and nucleoplasm-enriched genes (Supplementary Figure 1B), we observed a lower splicing efficiency on the transcripts from chromatin-enriched genes. We confirmed the HeLa results with the Raji chromatin RNA-seq and total RNA-seq datasets (Supplementary Figure 5C). We show as examples *PKM* and *NFIA*, a nucleoplasm-enriched and a chromatin-enriched gene, respectively, with co-transcriptional splicing events indicated by a star (Figure 4D). While co-transcriptional splicing is visible on both *NFIA* and *PKM, NFIA* has stronger intronic signals compared to *PKM*, especially on the chromatin RNA-seq and nucleoplasm RNA-seq. We also investigated whether there is a more general correlation between splicing efficiency in the POINT-seq or RNA-seq data and the production of mature mRNAs (nucleoplasm-enrichment ratio) (Figure 4E). Interestingly, we find that co-transcriptional splicing efficiency of nascent RNA does not correlate as well as the splicing efficiency of chromatin and nucleoplasm RNA-seq with the nucleoplasm-enrichment ratio (R=0.25 for POINT-seq vs R=0.38-0.42 for RNA-seq) (Figure 4E).

**Figure 4.**
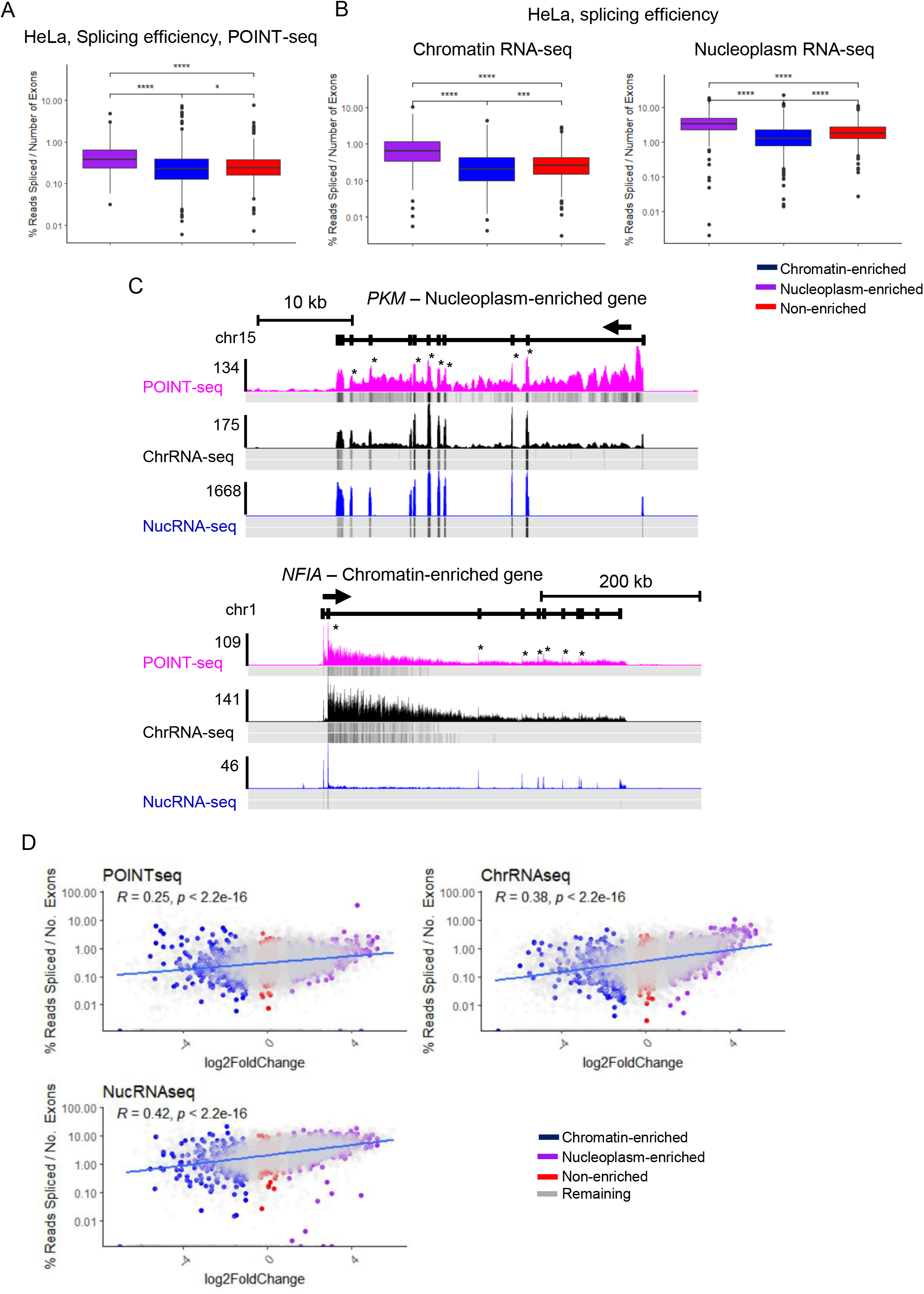
Transcripts from chromatin-enriched genes are less co-transcriptionally spliced. **(A)** Boxplots, shown as min to max with first quartile, median, and third quartile, of the splicing index of each transcript normalized to the number of exons from the POINT-seq data in HeLa cells of the chromatin RNA-seq 500 nucleoplasm-enriched (purple), chromatin-enriched (blue), or non-enriched (red) genes. Statistical test: Wilcoxon rank sum test. P-value: * < 0.05, **** < 0.0001. **(B)** Boxplots, shown as min to max with first quartile, median, and third quartile, of the splicing index of each transcript normalized to the number of exons from the chromatin RNA-seq and nucleoplasm RNA-seq data in HeLa of the chromatin RNA-seq 500 nucleoplasm-enriched (purple), chromatin-enriched (blue), or non-enriched (red) genes. Statistical test: Wilcoxon rank sum test. P-value: **** < 0.0001. **(C)** Screenshot of the genome browser POINT-seq, chromatin RNA-seq, and nucleoplasm RNA-seq tracks of the protein-coding genes *PKM* (nucleoplasm-enriched) and *NFIA* (chromatin-enriched). The read density of one biological replicate for each ChIP-seq is shown in colour while the other biological replicates density is shown below. Stars indicate the location of co-transcriptional splicing events. The arrow indicates the sense of transcription. **(D)** XY correlation plots of the nucleoplasm fold enrichment, defined as the fold change between nucleoplasm RNA-seq versus chromatin RNA-seq, and of the splicing index of each transcript normalized to the number of exons from POINT-seq, chromatin RNA-seq, or nucleoplasm RNA-seq. The Pearson correlation with p-value is indicated on each plot. Nucleoplasm-enriched (purple), chromatin-enriched (blue), or non-enriched (red), and remaining (grey) genes are shown.

As co-transcriptional splicing is associated with deposition of trimethylation on histone H3 lysine 36 (H3K36me3) by SETD2 (53), we re-analysed HeLa mNuc-seq datasets for H3K36me3 (54) (Supplementary Figure 5D and 5E). Interestingly, chromatin-enriched genes have lower H3K36me3 across the gene body, in line with less efficient co-transcriptional splicing.

As poor co-transcriptional splicing is also associated with transcriptional readthrough due to failure to recognise the poly(A) site (15,16), we analysed HeLa cell ChIP-seq of three CPA factors, CPSF73, PCF11, and Xrn2 (Figure 5A) (29). The nucleoplasm-enriched genes have clear peaks of CPA factors around the poly(A) site while the chromatin-enriched genes do not, suggesting inefficient mRNA CPA, as shown on *PKM* and *NFIA* (Figure 5B). To investigate mRNA CPA efficiency, we calculated the read-through index (RTI), which measures pol II pausing downstream of the poly(A) site (40), from Ser2P mNET-seq and observed that chromatin-enriched genes have higher transcriptional readthrough compared to the nucleoplasm-enriched and unchanged genes (Figure 5C and D and Supplementary Figure 6A). We confirmed this observation by re-analysing mNET-seq and chromatin RNA-seq data treated with siLuc or siCPSF73 (Figure 5E and F and Supplementary Figure 6B and 6C) (11,40). The chromatin-enriched genes are less sensitive to the loss of CPSF73 compared to non-enriched and nucleoplasm-enriched genes, which have more transcriptional readthrough following siCPSF73 treatment.

**Figure 5.**
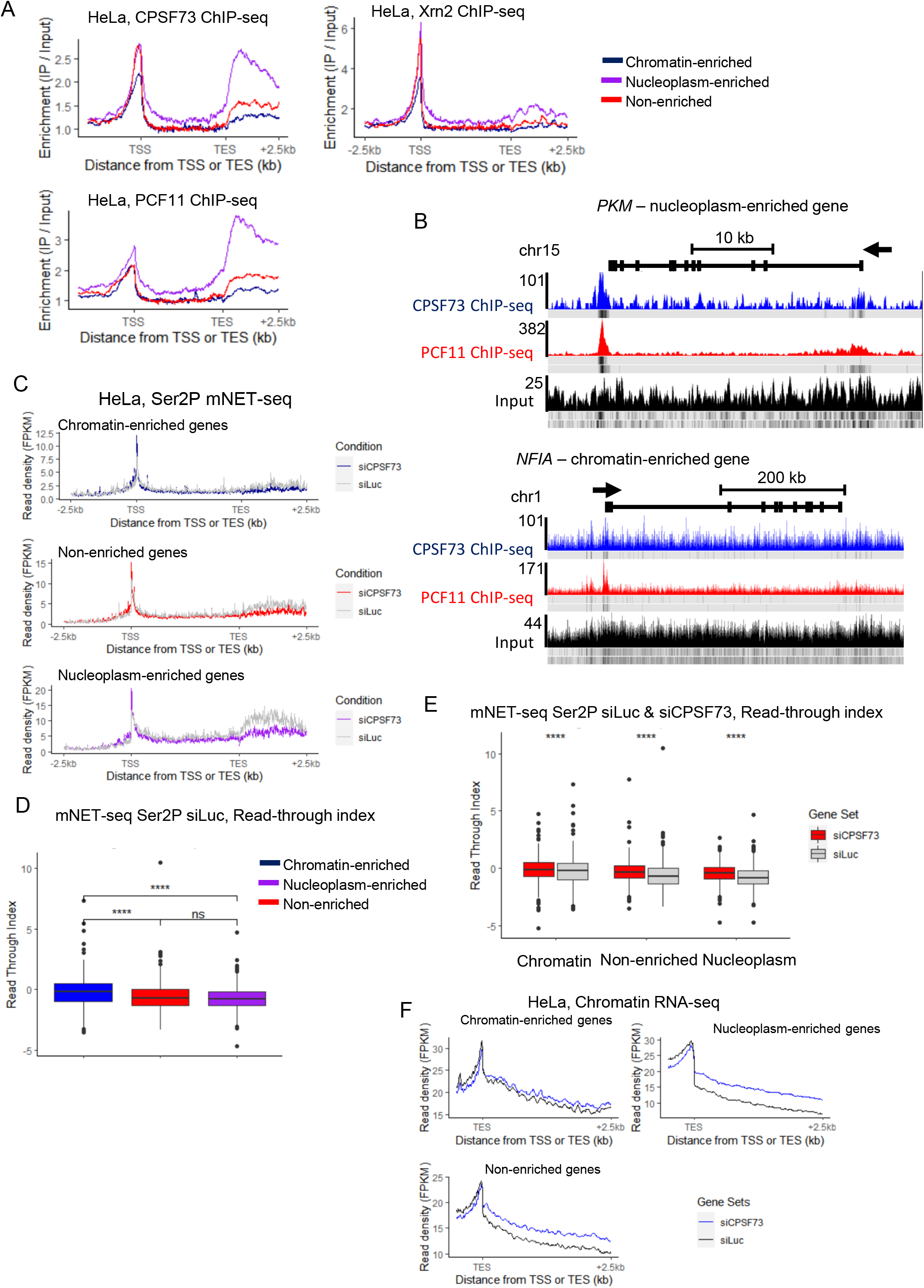
Transcripts from chromatin-enriched genes have a weak mRNA cleavage and polyadenylation. Metagene profiles in HeLa cells of CPSF73, PCF11, and Xrn2 ChIP-seq of the chromatin RNA-seq 500 nucleoplasm-enriched (purple), chromatin-enriched (blue), or non-enriched (red) genes. **(B)** Screenshot of the genome browser ChIP-seq tracks of the protein-coding genes *PKM* (nucleoplasm-enriched) and *NFIA* (chromatin-enriched). The read density of one biological replicate for each ChIP-seq is shown in colour while the other biological replicates density is shown below. The arrow indicates the sense of transcription. **(C)** Metagene profiles in HeLa cells of Ser2P mNET-seq treated with siLuc (grey) or siCPSF73 (coloured) of the chromatin RNA-seq 500 nucleoplasm-enriched, chromatin-enriched, or non-enriched genes. **(D)** Boxplots, shown as min to max with first quartile, median, and third quartile, of the read-through index calculated on the Ser2P mNET-seq treated with siLuc of the chromatin RNA-seq 500 nucleoplasm-enriched (purple), chromatin-enriched (blue), or non-enriched (red) genes. Statistical test: Wilcoxon rank sum test. P-value: ns: not significant, **** < 0.0001. **(E)** Boxplots, shown as min to max with first quartile, median, and third quartile, of the read-through index calculated on the Ser2P mNET-seq treated with siLuc (grey) or siCPSF73 (red) of the chromatin RNA-seq 500 nucleoplasm-enriched, chromatin-enriched, or non-enriched genes. Statistical test: Wilcoxon rank sum test. P-value: **** < 0.0001. **(F)** Metagene profiles in HeLa cells of chromatin RNA-seq treated with siLuc (black) or siCPSF73 (blue) of the chromatin RNA-seq 500 nucleoplasm-enriched, chromatin-enriched, or non-enriched genes.

These findings demonstrate that even though chromatin-enriched gene transcripts are on average longer, these transcripts are poorly processed, which could explain their chromatin retention.

### Transcripts from chromatin-enriched genes are sensitive to the nuclear RNA exosome

The nuclear RNA exosome complex promotes RNA degradation, for example of pre-mRNAs with processing defects, such as those with retained introns or transcriptional readthrough (55). We investigated whether the nuclear RNA exosome complex could degrade the chromatin-enriched gene transcripts, which are poorly processed. We re-analysed previously published HeLa nucleoplasm RNA-seq treated with siLuc or siEXOSC3 (siEX3), a core component of the nuclear RNA exosome activity (Supplementary Figure 7A) (11). 551 transcripts, including 518 from protein-coding genes were downregulated and 1,926 transcripts, including 427 from protein-coding genes, were up-regulated after depletion of the RNA exosome. Comparison of siEX3 upregulated genes (siEX3(+)) with chromatin-enriched or nucleoplasm-enriched genes shows only a moderate correlation (R = -0.29, p < 2.2e-16) (Supplementary Figure 7B). This initial analysis of the nuclear RNA exosome knockdown shows a limited effect on the mRNA levels based on exons. To determine whether an effect is observed on intron retention, a splicing defect targeted by the nuclear RNA exosome (55), we used rMATS on the chromatin and nucleoplasm RNA-seq before and after siEXOSC3 to obtain the list of significant splicing changes, including alternative 5’ and 3’ splice sites, mutually exclusive exons, retained introns, and skipped exons (Figure 6A and Supplementary Figure 7C). We find a specific increase in intron retention cases in nucleoplasm RNA-seq following siEXOSC3, indicating that these poorly-processed transcripts are usually degraded by the nuclear RNA exosome. To determine whether transcripts targeted by the nuclear RNA exosome are coming from chromatin-enriched, non-enriched, or nucleoplasm-enriched genes, we compared the changes in expression across the whole transcript units (exons and introns) before and after treatment with siEXOSC3 (Figure 6B and Supplementary Figure 7D and 7E). We found that the expression of transcripts from chromatin-enriched genes is more increased after siEXOSC3 compared to transcripts from non-enriched and nucleoplasm-enriched genes. To confirm the observation that the increase of transcripts from chromatin-enriched genes are also poorly processed, we calculated the splicing efficiency before and after siEXOSC3 (Figure 6C and Supplementary Figure 7F). Importantly, we found that the splicing efficiency of transcripts from chromatin-enriched genes are the most decreased following knockdown of the nuclear RNA exosome, which indicates an increase in the chromatin and nucleoplasm of poorly-processed transcripts from these genes. The increase in poorly-spliced transcripts from chromatin-enriched genes after siEXOSC3 can be observed on single gene examples, *NFIA* and *MDM4*, while no obvious changes in RNA splicing are visible for the transcripts of the nucleoplasm-enriched genes *PKM* and *PSAP* (Figure 6D).

**Figure 6.**
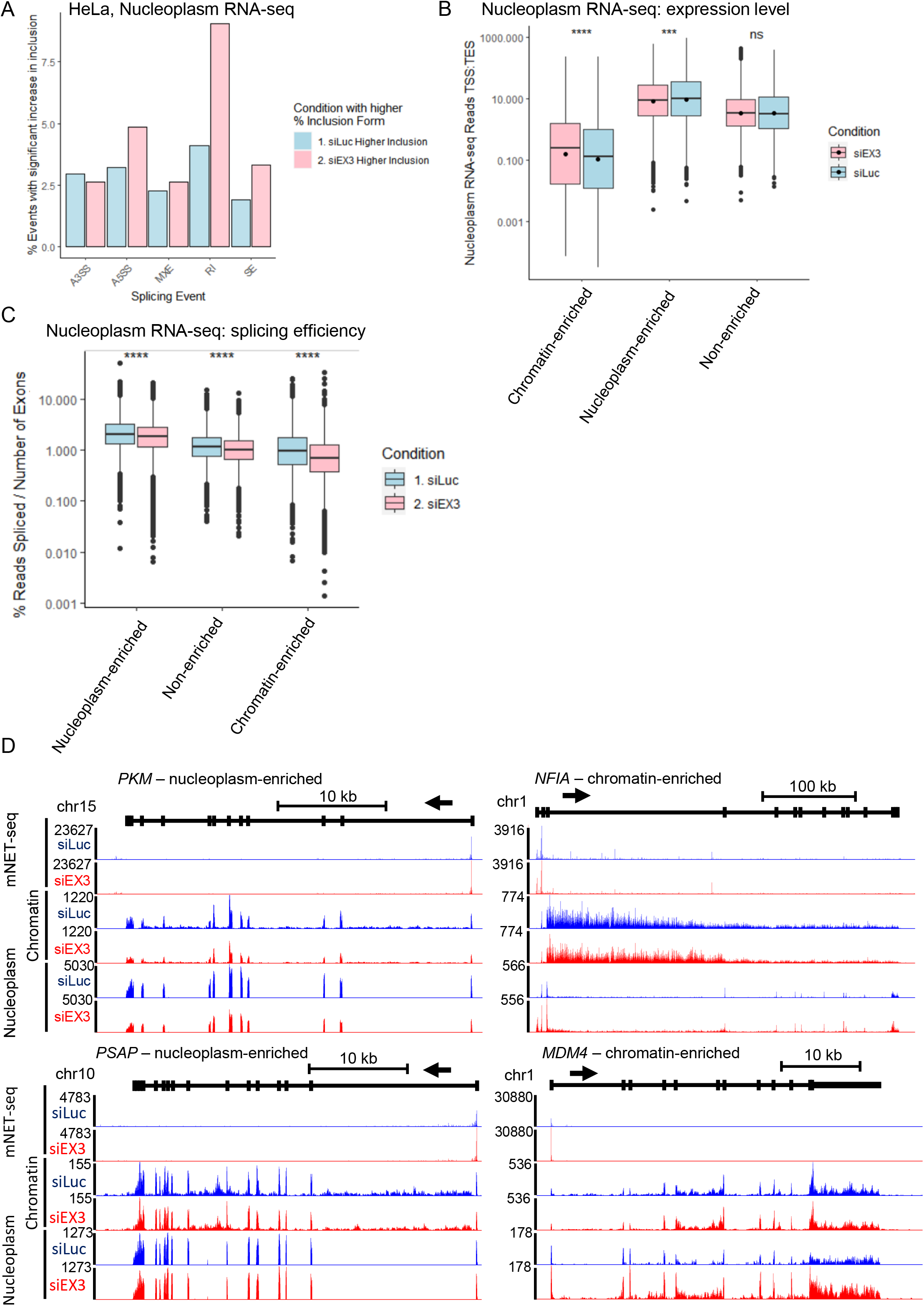
Transcripts from chromatin-enriched genes are sensitive to the nuclear RNA exosome. **(A)** Bar charts of significant changes in nucleoplasm RNA-seq of splicing events obtained with rMATs in control (siLuc, blue) or following the knockdown of the nuclear RNA exosome (siEX3, pink). A3SS: alternative 3’ splice site; A5SS: alternative 5’ splice site, MXE: mutually exclusive exons; IR: intron retention; SE: skipped exon. **(B)** Boxplots, shown as min to max with first quartile, median, and third quartile, of the expression level of full-length transcripts, including exons and introns, from the nucleoplasm RNA-seq data in HeLa cells in control (siLuc, blue) or after siEXOSC3 knockdown (siEX3, pink) of all nucleoplasm-enriched, chromatin-enriched, or non-enriched genes. Statistical test: Wilcoxon rank sum test. P-value: n.s. not significant, *** < 0.001, **** < 0.0001. **(C)** Boxplots, shown as min to max with first quartile, median, and third quartile, of the splicing index of each transcript normalized to the number of exons from the nucleoplasm RNA-seq data in HeLa cells in control (siLuc, blue) or after siEXOSC3 knockdown (siEX3, pink) of all nucleoplasm-enriched, chromatin-enriched, or non-enriched genes. Statistical test: Wilcoxon rank sum test. P-value: n.s. not significant, **** < 0.0001. **(D)** Screenshot of the genome browser total pol II mNET-seq, chromatin RNA-seq, and nucleoplasm RNA-seq tracks treated with siLuc (blue) or siEXOSC3 (red) of the protein-coding genes *PKM* and *PSAP* (nucleoplasm-enriched) and *NFIA* and *MDM4* (chromatin-enriched). The arrow indicates the sense of transcription.

These findings indicate that poorly-processed transcripts from chromatin-enriched genes that are released from the chromatin are degraded by the nuclear RNA exosome.

## DISCUSSION

Production of mature mRNA requires both transcription and co/post-transcriptional RNA processing. Regulation of gene expression via the control of transcription initiation and pol II pause release are well established (56). We show here that the efficiency of RNA processing is also an important factor controlling chromatin retention of transcripts, potentially because of higher R-loop levels (57), and the degradation in the nucleoplasm by the nuclear RNA exosome of poorly-processed transcripts (55,58). There is a large subgroup of protein-coding genes that are transcribed but the transcripts are poorly processed and chromatin retained, which results in poor production of mature mRNAs and proteins. Interestingly, we found that this subset of chromatin-enriched genes shares transcriptional and co-transcriptional similarities with lncRNA genes (11). These include higher CTD Thr4 phosphorylation, poor pre-mRNA splicing and CPA, higher transcriptional readthrough, decreased sensitivity to CPSF73 KD, higher chromatin retention, and degradation of the poorly-processed transcripts in the nucleoplasm by the nuclear RNA exosome (Figure 7). Together these observations explain the low levels of mature mRNAs from these genes and indicate that the cellular mechanisms that regulate levels of lncRNAs are also used to regulate expression of protein-coding genes. Of interest, Schlackow et al (11) also found a small subset of lncRNA genes whose transcripts are processed efficiently and not retained on the chromatin. These observations indicate that there is overlap between protein-coding genes and lncRNA genes in terms of the mechanisms operating at the transcriptional and co-transcriptional levels. The efficiency of transcription and co-transcriptional processes across transcription units, including protein-coding and non-coding genes, can be viewed as a continuum with poorly-expressed and poorly-processed lncRNAs at one end and highly-expressed and efficiently processed mRNAs at the other end with some overlap in the middle.

**Figure 7.**
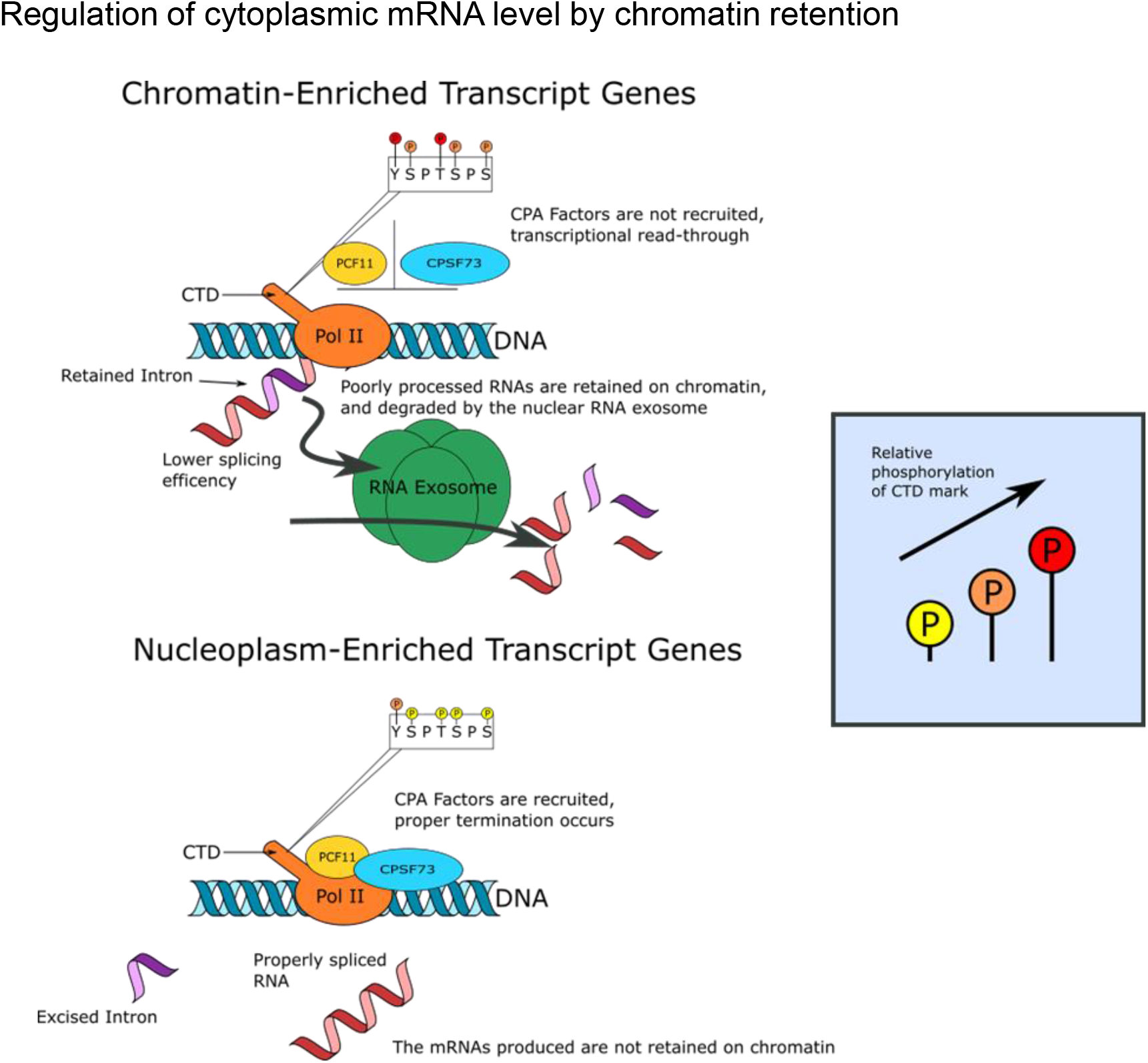
RNA processing efficiency regulates cytoplasmic mRNA level via a combination of chromatin retention and nuclear RNA degradation. The high production of mature mRNAs of nucleoplasm-enriched genes is associated with a more efficient pre-mRNA splicing and mRNA CPA, resulting in the production of more stable mRNA that will be exported to the cytoplasm to be translated. In contrast, transcripts from chromatin-enriched genes are associated with higher phosphorylation of pol II CTD Thr4 residues, less efficient pre-mRNA splicing and mRNA CPA, chromatin retention of the poorly processed transcripts, and a shorter mRNA half-life due to degradation by the nuclear RNA exosome of the poorly processed transcripts that are located in the nucleoplasm.

Some of the chromatin-enriched genes are well transcribed but produce hardly any mature mRNAs and proteins, which begs the question: what is the cellular advantage of transcribing a protein-coding gene without producing a protein? It is possible that these genes are transcribed but the transcripts are poorly processed until their proteins are required, which would require only the activation of RNA processing. Our data indicate that the downregulation of these genes occurs via poor co-transcriptional RNA processing, chromatin retention, and degradation of the transcripts by the nuclear RNA exosome rather than low transcription. In addition, overlap between the HeLa and Raji datasets show a higher proportion of common genes between nucleoplasm-enriched genes (47%-63%) compared to chromatin-enriched genes (19%-44%), which indicates a higher diversity in transcribed but poorly-processed transcript genes, at least between these two cancer cell lines. Interestingly, while we found only ∼160 genes that are in opposing categories (chromatin-enriched in one cell line and nucleoplasm-enriched in the other, or vice-versa) between the two cell lines, this indicates that genes could potentially move from one category to another depending on the cell line or following a cellular stress, for example.

A surprising observation is the lower CTD phosphorylation level on the nucleoplasm-enriched genes, especially the genes with the highest nucleoplasm-enrichment ratios. As pol II CTD phosphorylation is known to recruit splicing proteins and mRNA CPA factors (59), it is unexpected that lower Ser2P and Ser5P levels are associated with better RNA processing. Slow pol II elongation has been shown to result in hyperphosphorylation of the CTD Ser2 residues at the 5’ end of genes, promoting a higher dwell time at start sites and a reduced transcriptional polarity (60). In addition, hyperphosphorylation of the CTD during M phase inhibits pol II, which contributes to mitotic gene silencing (61,62). While more work is required, these observations suggest that the level of pol II CTD phosphorylation could play an important role in controlling transcription activity and co-transcriptional processing efficiency. However, the decrease in CTD phosphorylation for the nucleoplasm-enriched genes or the higher Tyr1 and Thr4 phosphorylation for chromatin-enriched genes was generally observed only after normalisation to total pol II. The measurement of pol II CTD phosphorylation using antibody-based technique contains several potential pitfalls, including different affinities of each antibody, the influence of other pol II CTD modifications on the antibody specificity, and CTD-interacting proteins that can influence antibody accessibility.

We previously found that inhibition of the protein phosphatase PP2A causes a higher production of poly(A)+ mRNA without any significant changes in transcription level (34), and this is likely due to more efficient cleavage and polyadenylation. Modulation of RNA processing efficiency can therefore regulate gene expression at several different levels.

## SUPPLEMENTARY DATA

Supplementary Data are available at NAR online.

## ACKNOWLEDGEMENT

We thank Dr Andrew Angel (University of Aberdeen, UK) and Prof Nick. J Proudfoot (University of Oxford, UK) for discussion.

## Author contributions

C.H carried out all the bioinformatics analysis with supervision from M.T. S.M. supervised C.H. and M.T. M.T. and S.M. designed the project. M.T. wrote the paper with contributions from all the authors.

