## Supplementaty Figures for "Cytoplasmic mRNA levels are regulated by a combination of chromatin retention and RNA stability"

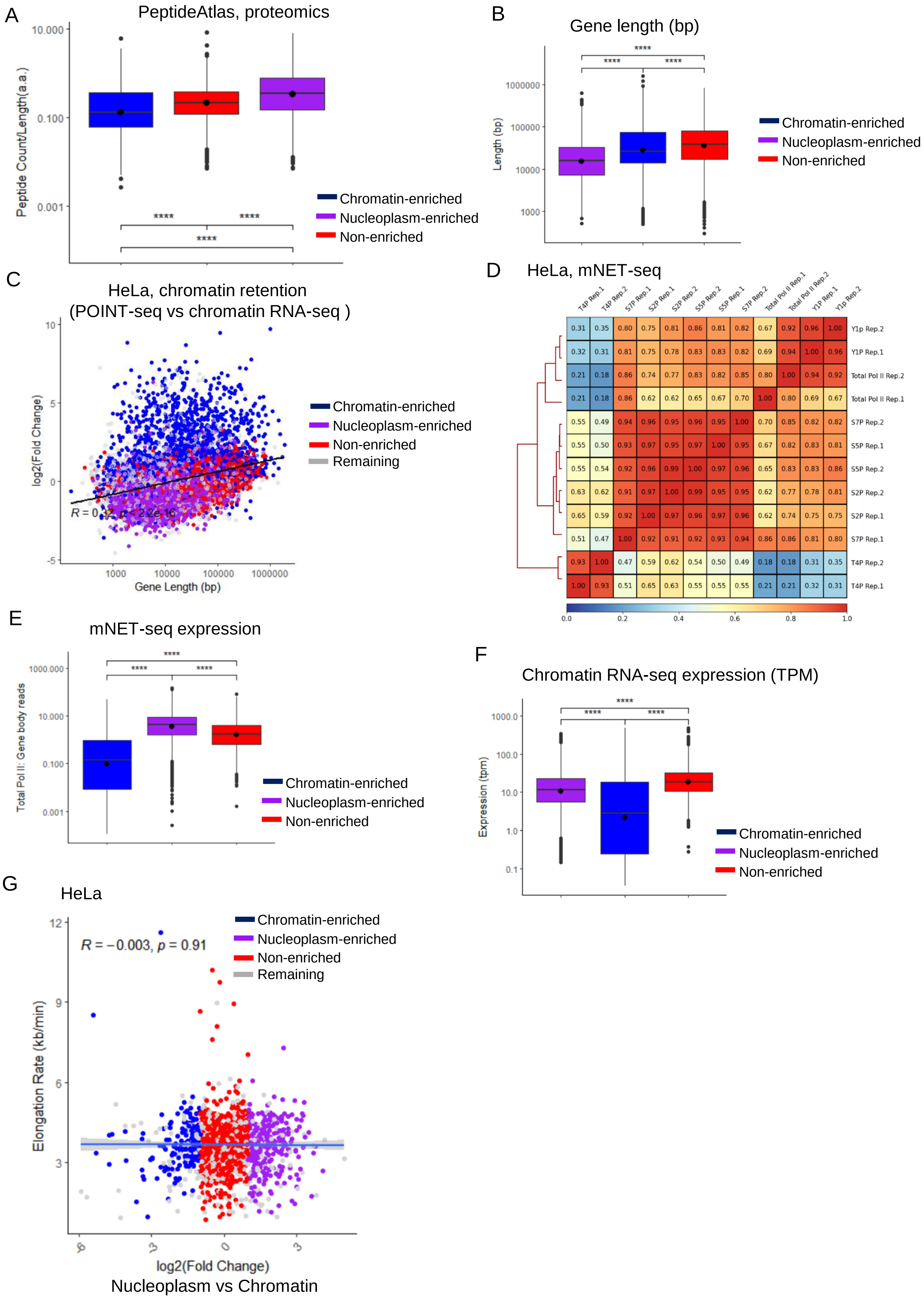


Supplementary Figure 1

**Supplementary Figure 1. Higher levels of pol II Tyr1 and Thr4 phosphorylation are associated with poor expression and chromatin retention of transcripts**.

**(A)** Boxplots, shown as min to max with first quartile, median, and third quartile, of the number of peptides per protein found in the PeptideAtlas database for the nucleoplasm-enriched (purple), chromatin-enriched (blue), or non-enriched (red) genes. The number of proteins found to have at least one peptide are indicated at the top of each category. Statistical test: Wilcoxon rank sum test. P-value: **** < 0.0001. **(B)** Correlation matrix heatmap for the mNET-seq showing positive correlation in red and absence of correlation in blue. **(C)** Boxplots, shown as min to max with first quartile, median, and third quartile, of the expression in total pol II mNET-seq of the nucleoplasm-enriched (purple), chromatin-enriched (blue), or non-enriched (red) genes. **(D)** Boxplots, shown as min to max with first quartile, median, and third quartile, of the expression (TPM) in chromatin RNA-seq of the nucleoplasm-enriched (purple), chromatin-enriched (blue), or non-enriched (red) genes. **(E)** Boxplots, shown as min to max with first quartile, median, and third quartile, of the gene length of the nucleoplasm-enriched (purple), chromatin-enriched (blue), or non-enriched (red) genes. **(F)** XY correlation plot of the gene length (bp) versus the log2 fold change of “Chromatin RNA-seq versus POINT-seq”, which provides a measure of chromatin retention. The Pearson correlation with p-value is indicated on the plot. Nucleoplasm-enriched (purple), chromatin-enriched (blue), non-enriched (red), and remaining (grey) genes are shown. **(G)** Screenshot of the genome browser chromatin RNA-seq, nucleoplasm RNA-seq, cytoplasm RNA-seq, and total pol II, Tyr1P, Ser2P, Thr4P, Ser5P, and Ser7P mNET-seq tracks of the protein-coding genes *PKM* (nucleoplasm-enriched) and *NFIA* (chromatin-enriched). The arrow indicates the sense of transcription.

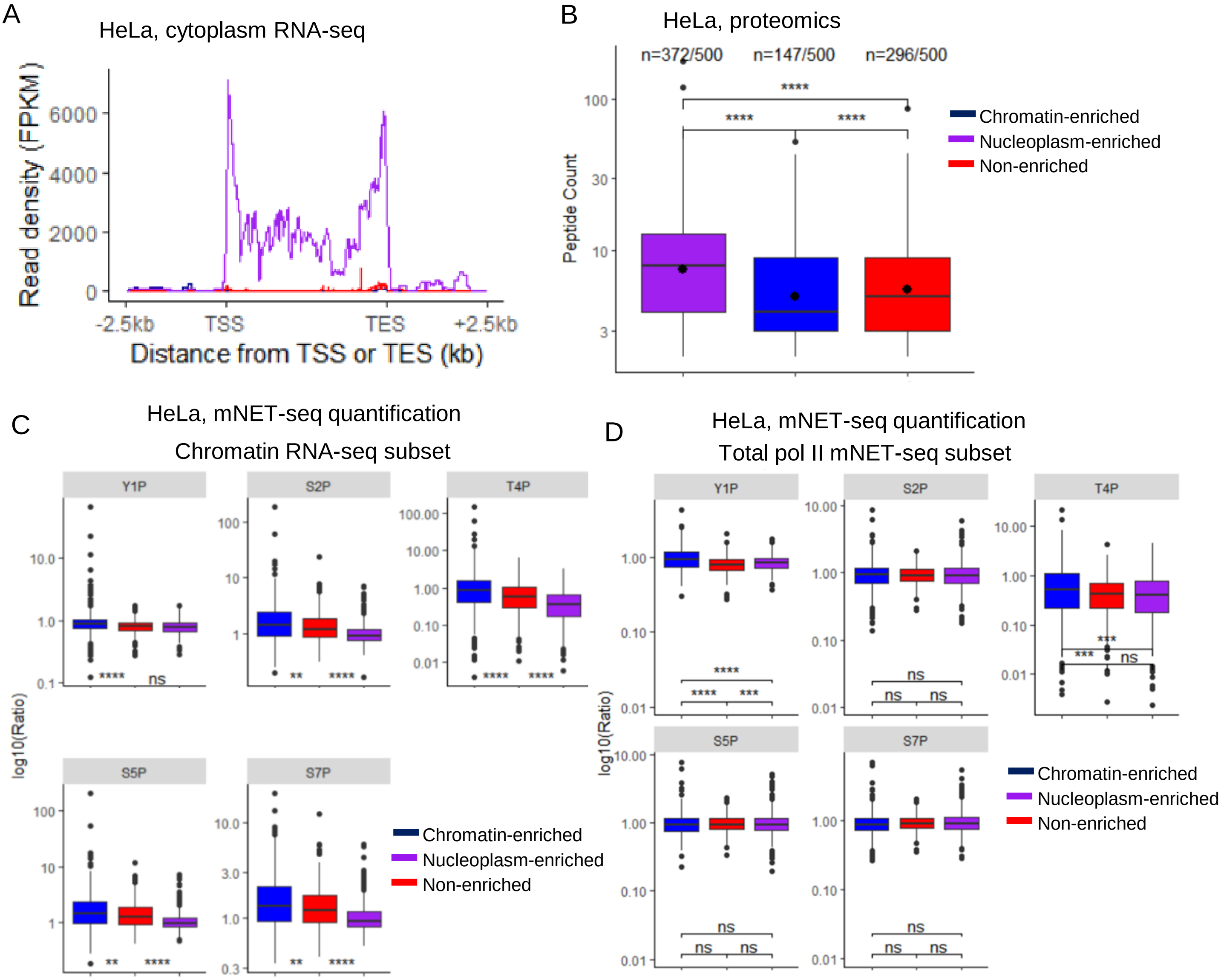


Supplementary Figure 2

**Supplementary Figure 2. Transcripts from chromatin-enriched genes are associated with a higher phosphorylation of the pol II CTD Thr4 residues.**

**(A)** Metagene profiles in HeLa cells of cytoplasm RNA-seq of the 500 nucleoplasm-enriched (purple), chromatin-enriched (blue), or non-enriched (red) genes. **(B)** Boxplots, shown as min to max with first quartile, median, and third quartile, of the number of peptides per protein found for the 500 nucleoplasm-enriched (purple), chromatin-enriched (blue), or non-enriched (red) genes. The number of proteins found to have at least one peptide are indicated at the top of each category. Statistical test: Wilcoxon rank sum test. P-value: **** < 0.0001. **(C)** Boxplots, shown as min to max with first quartile, median, and third quartile, of each pol II CTD phosphorylation mark ratioed to total pol II across the gene body of the chromatin RNA-seq selected 500 nucleoplasm-enriched (purple), chromatin-enriched (blue), and non-enriched (red) genes. Statistical test: Wilcoxon rank sum test. P-value: ns: not significant, ** < 0.01, **** < 0.0001. **(D)** Boxplots, shown as min to max with first quartile, median, and third quartile, of each pol II CTD phosphorylation mark ratioed to total pol II across the gene body of the total pol II mNET-seq selected nucleoplasm-enriched (purple), chromatin-enriched (blue), and non-enriched (red) genes. Statistical test: Wilcoxon rank sum test. P-value: ns: not significant, *** < 0.001, **** < 0.0001.

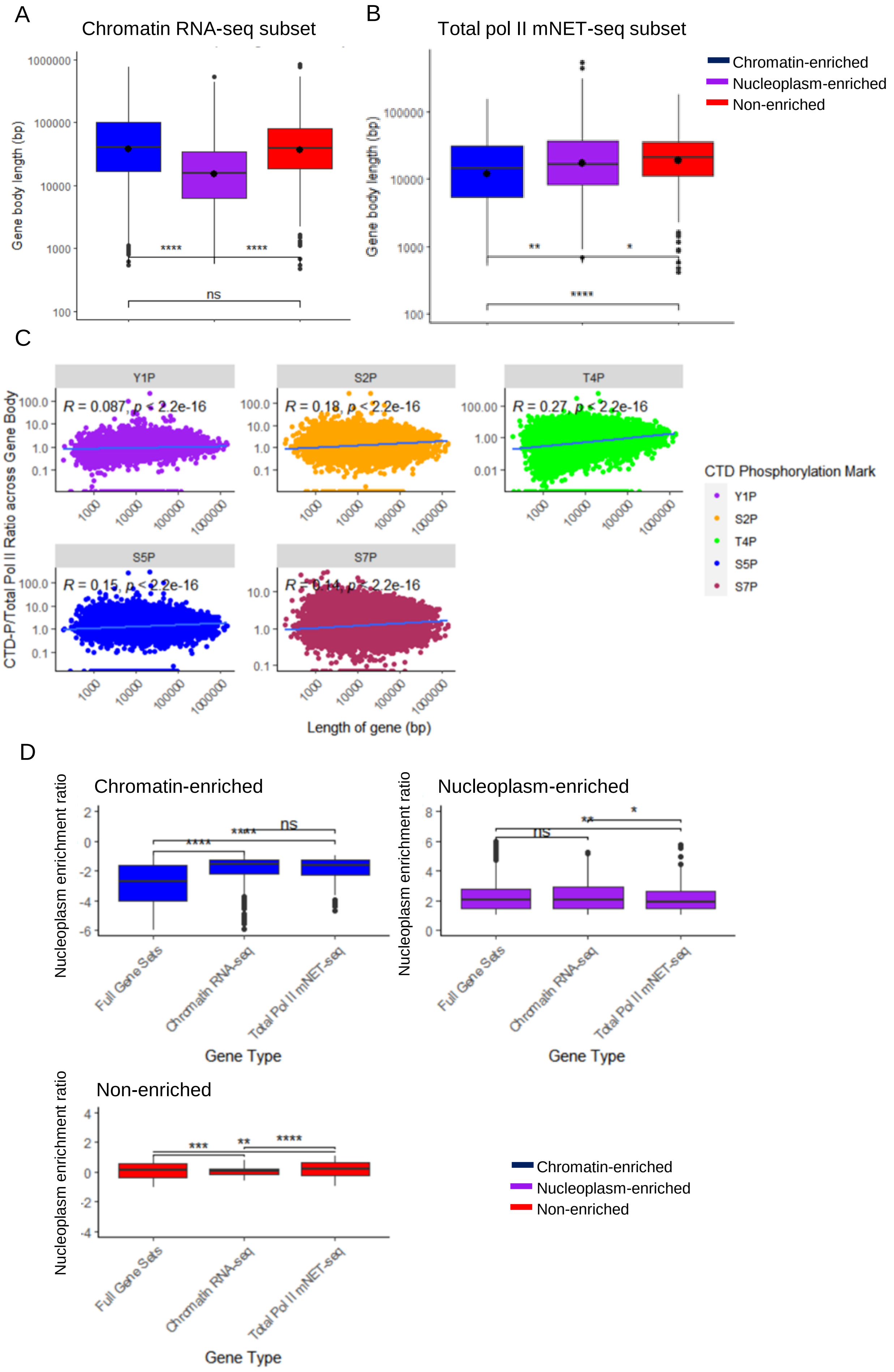


Supplementary Figure 3

**Supplementary Figure 3. Genes expressing the highest nucleoplasm-enriched transcripts have less .**

**(A)** Boxplots, shown as min to max with first quartile, median, and third quartile, of the gene length of the 500 chromatin RNA-seq selected nucleoplasm-enriched (purple), chromatin-enriched (blue), and non-enriched (red) genes. Statistical test: Wilcoxon rank sum test. P-value: n.s. not significant, **** < 0.0001. **(B)** Boxplots, shown as min to max with first quartile, median, and third quartile, of the gene length of the 10% total pol II mNET-seq selected nucleoplasm-enriched (purple), chromatin-enriched (blue), and non-enriched (red) genes. Statistical test: Wilcoxon rank sum test. P-value: * < 0.05, ** < 0.01, **** < 0.0001. **(C)** XY correlation plots of the gene length (in bp) and each pol II CTD phosphorylation mark ratioed to total pol II. The Pearson correlation with p-value is indicated on each plot. Nucleoplasm-enriched (purple), chromatin-enriched (blue), non-enriched (red), and remaining (grey) genes are shown. **(D)** Boxplots, shown as min to max with first quartile, median, and third quartile, of the nucleoplasm-enrichment ratio (log2 fold change of nucleoplasm RNA-seq versus chromatin RNA-seq) of the total gene set, the 500 chromatin RNA-seq selected, and the 10% total pol II mNET-seq selected nucleoplasm-enriched (purple), chromatin-enriched (blue), and non-enriched (red) genes. Statistical test: Wilcoxon rank sum test. P-value: n.s. not significant, * < 0.05, ** < 0.01, *** < 0.001, **** < 0.0001.

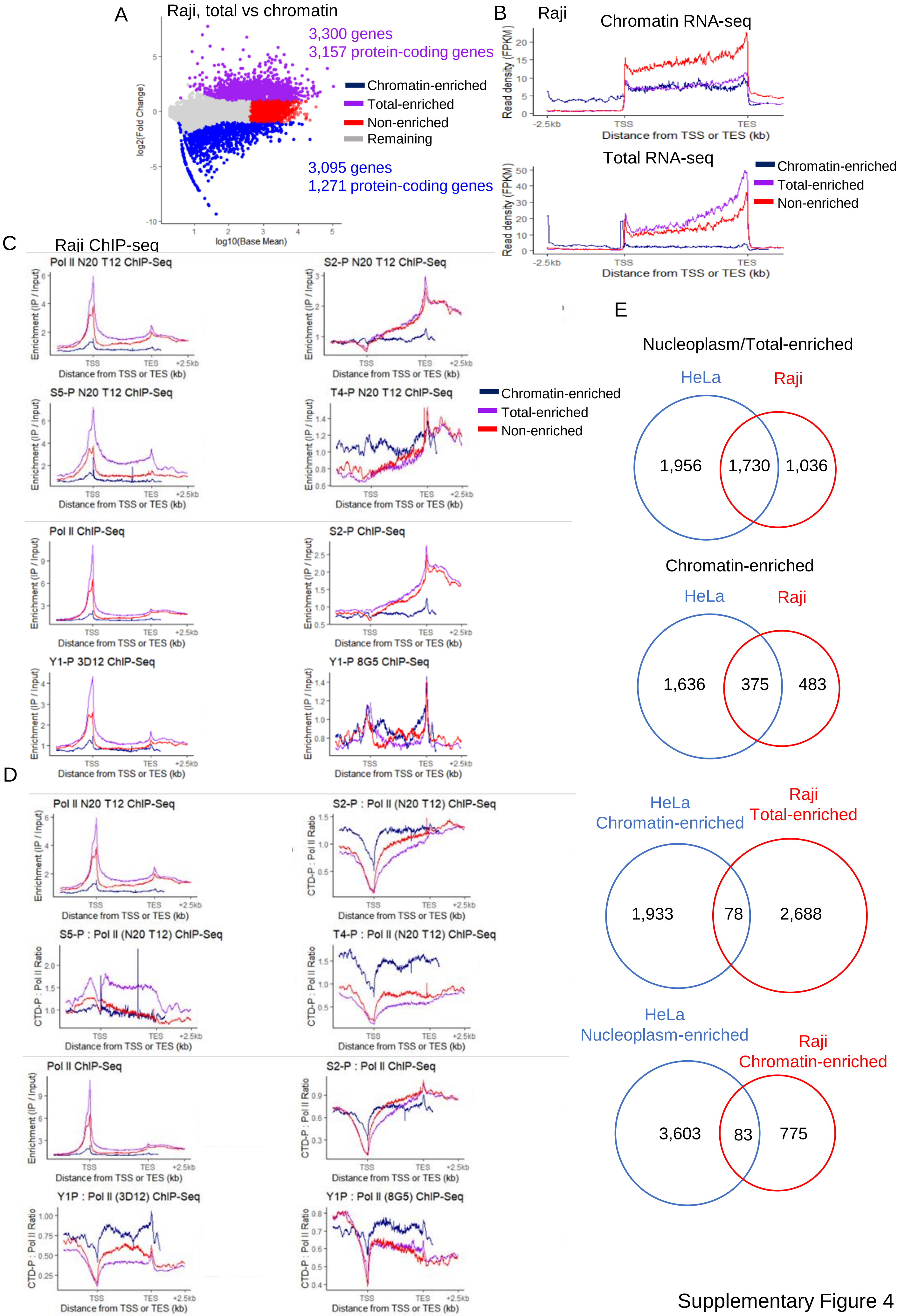


Supplementary Figure 4

**Supplementary Figure 4. Raji pol II CTD ChIP-seq show similar patterns to HeLa pol II CTD mNET-seq.**

**(A)** MA plot in Raji cells of the intron-containing protein-coding genes found to be differentially enriched in the total (total-enriched, purple) or in the chromatin (chromatin-enriched, blue) fraction. A set of non-enriched genes (red) and the remaining genes (grey) are also indicated. **(B)** Metagene profiles in Raji cells of chromatin RNA-seq and total RNA-seq of total-enriched (purple), chromatin-enriched (blue), or non-enriched (red) genes. **C)** Metagene profiles of total pol II, Tyr1P, Ser2P, Thr4P, and Ser5P in Raji cells across the Raji-defined total-enriched (purple), chromatin-enriched (blue), or non-enriched (red) genes. **(D)** Metagene profiles of each Tyr1P, Ser2P, Thr4P, or Ser5P signal ratioed to total pol II signal in Raji cells across the Raji-defined total-enriched (purple), chromatin-enriched (blue), or non-enriched (red) genes. **(E)** Overlap between nucleoplasm/total-enriched genes or between chromatin-enriched genes found to be expressed in both HeLa and Raji cells.

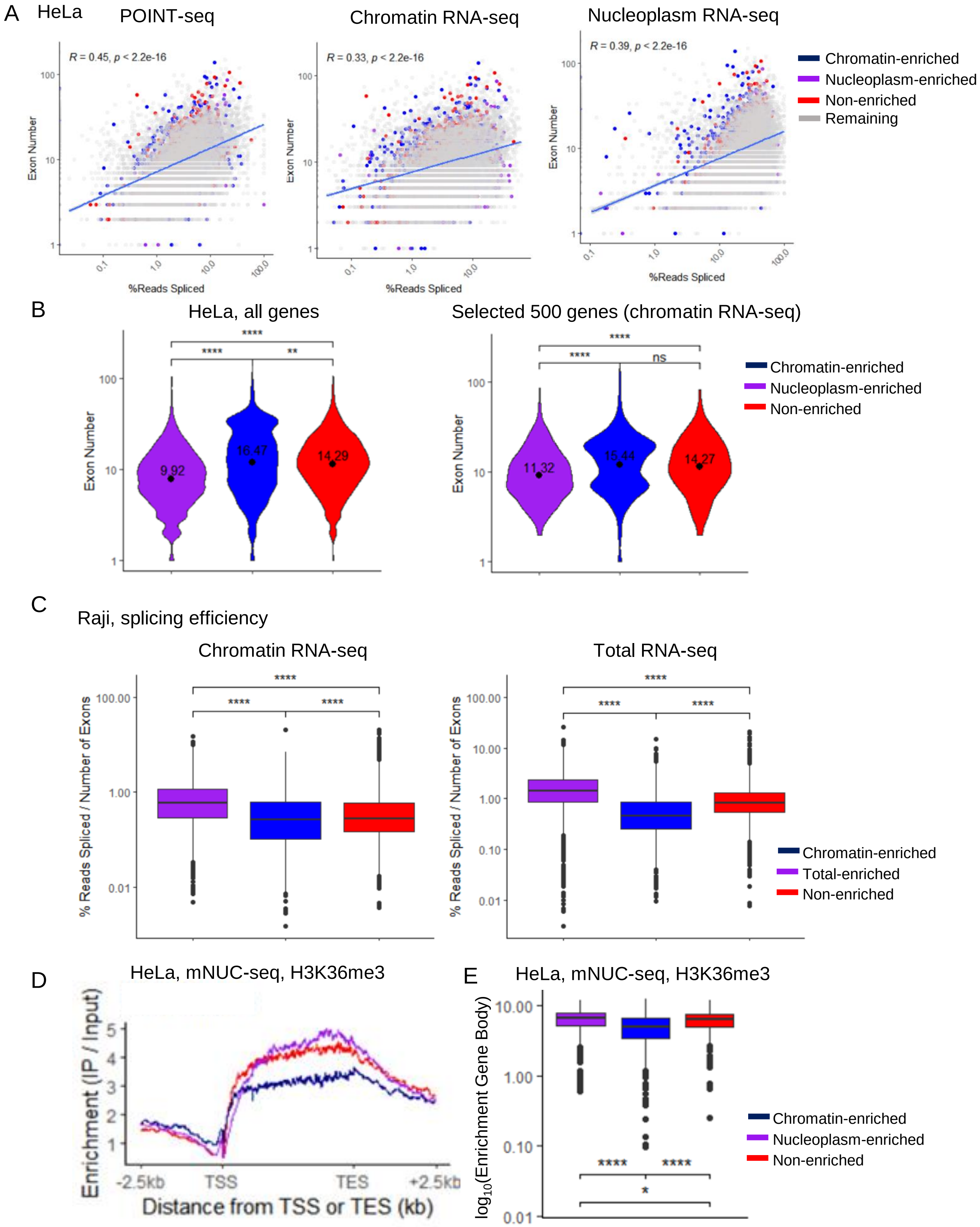


Supplementary Figure 5

**Supplementary Figure 5. Transcripts from chromatin-enriched genes are less co-transcriptionally spliced**.

**(A)** XY correlation plots of the splicing efficiency versus the number of exons of each transcript in the POINT-seq, chromatin RNA-seq, and nucleoplasm RNA-seq. The Pearson correlation with p-value is indicated on each plot. Nucleoplasm-enriched (purple), chromatin-enriched (blue), non-enriched (red), and remaining (grey) genes are shown. **(B)** Violin plots, shown as min to max with first quartile, median, and third quartile, of the number of exons in all, or in the chromatin RNA-seq selected 500, nucleoplasm-enriched (purple), chromatin-enriched (blue), non-enriched (red) genes. Statistical test: Wilcoxon rank sum test. P-value: ns: not significant, ** < 0.01, **** < 0.0001. **(C)** Metagene profiles in HeLa cells of H3K36me3 mNuc-seq of the nucleoplasm-enriched (purple), chromatin-enriched (blue), non-enriched (red) genes. H3K36me3 is shown across the whole gene body (TSS – 2.5 kb to TES + 2.5 kb). Metaprofiles are shown as IP / Input. **(D)** Boxplots, shown as min to max with first quartile, median, and third quartile, of each histone mark ratioed to Input of the nucleoplasm-enriched (purple), chromatin-enriched (blue), non-enriched (red) genes. Quantification is performed on reads located within TSS to TES. Statistical test: Wilcoxon rank sum test. P-value: ns: not significant, * < 0.05, ** < 0.01, **** < 0.0001. **(E)** Boxplots, shown as min to max with first quartile, median, and third quartile, of the splicing index of each transcript normalized to the number of exons from the chromatin RNA-seq and total RNA-seq data in Raji of the total-enriched (purple), chromatin-enriched (blue), non-enriched (red) genes. Statistical test: Wilcoxon rank sum test. P-value: **** < 0.0001.

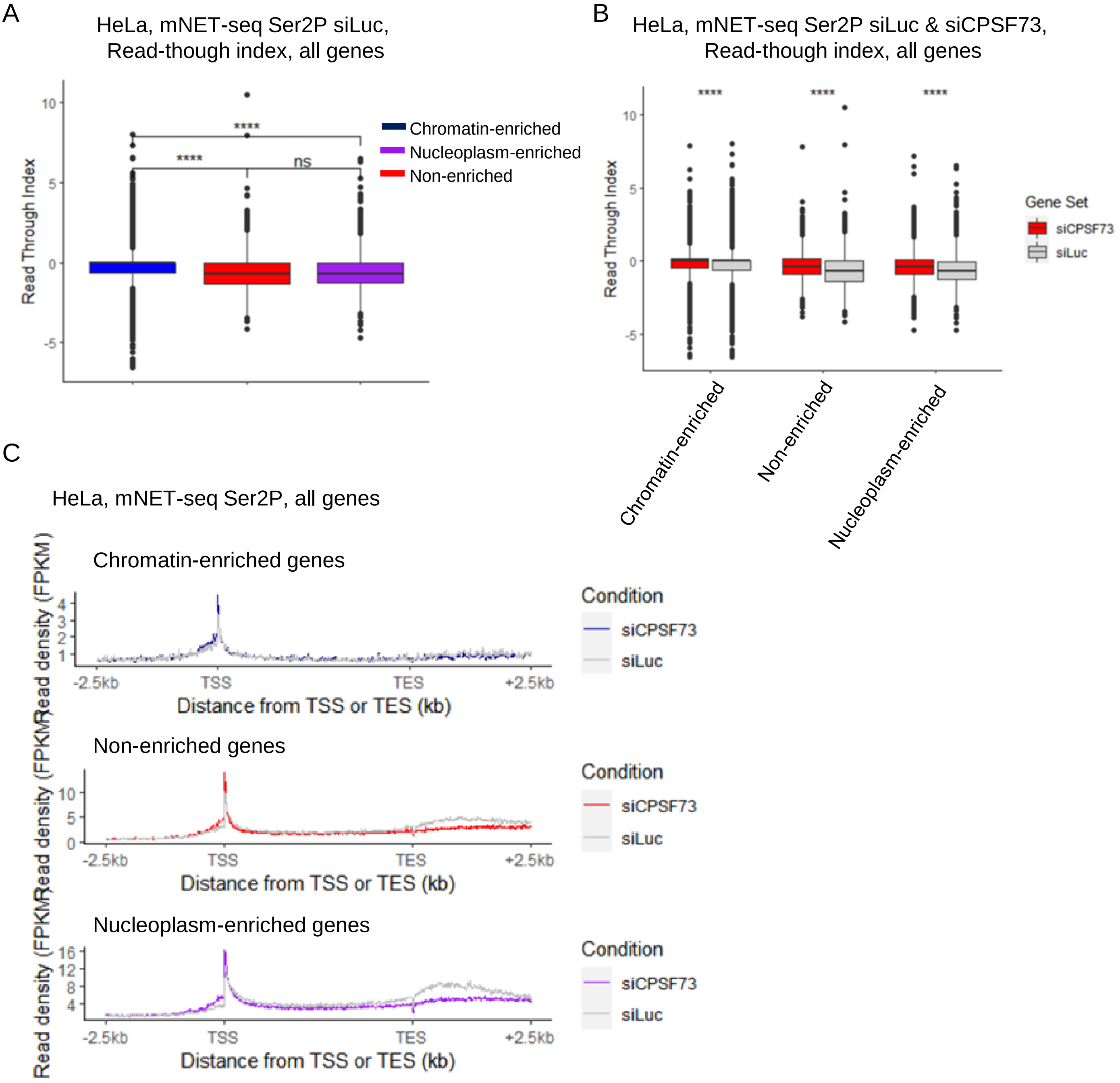


Supplementary Figure 6

**Supplementary Figure 6. Transcripts from chromatin-enriched genes have a weak mRNA cleavage and polyadenylation**.

**(A)** Boxplots, shown as min to max with first quartile, median, and third quartile, of the read-through index calculated on the Ser2P mNET-seq treated with siLuc of all nucleoplasm-enriched (purple), chromatin-enriched (blue), or non-enriched (red) genes. Statistical test: Wilcoxon rank sum test. P-value: ns: not significant, **** < 0.0001. **(B)** Boxplots, shown as min to max with first quartile, median, and third quartile, of the read-through index calculated on the Ser2P mNET-seq treated with siLuc (grey) or siCPSF73 (red) of all nucleoplasm-enriched, chromatin-enriched, or non-enriched genes. Statistical test: Wilcoxon rank sum test. P-value: **** < 0.0001. **(C)** Metagene profiles in HeLa cells of chromatin RNA-seq treated with siLuc (black) or siCPSF73 (blue) of all nucleoplasm-enriched, chromatin-enriched, or non-enriched genes.

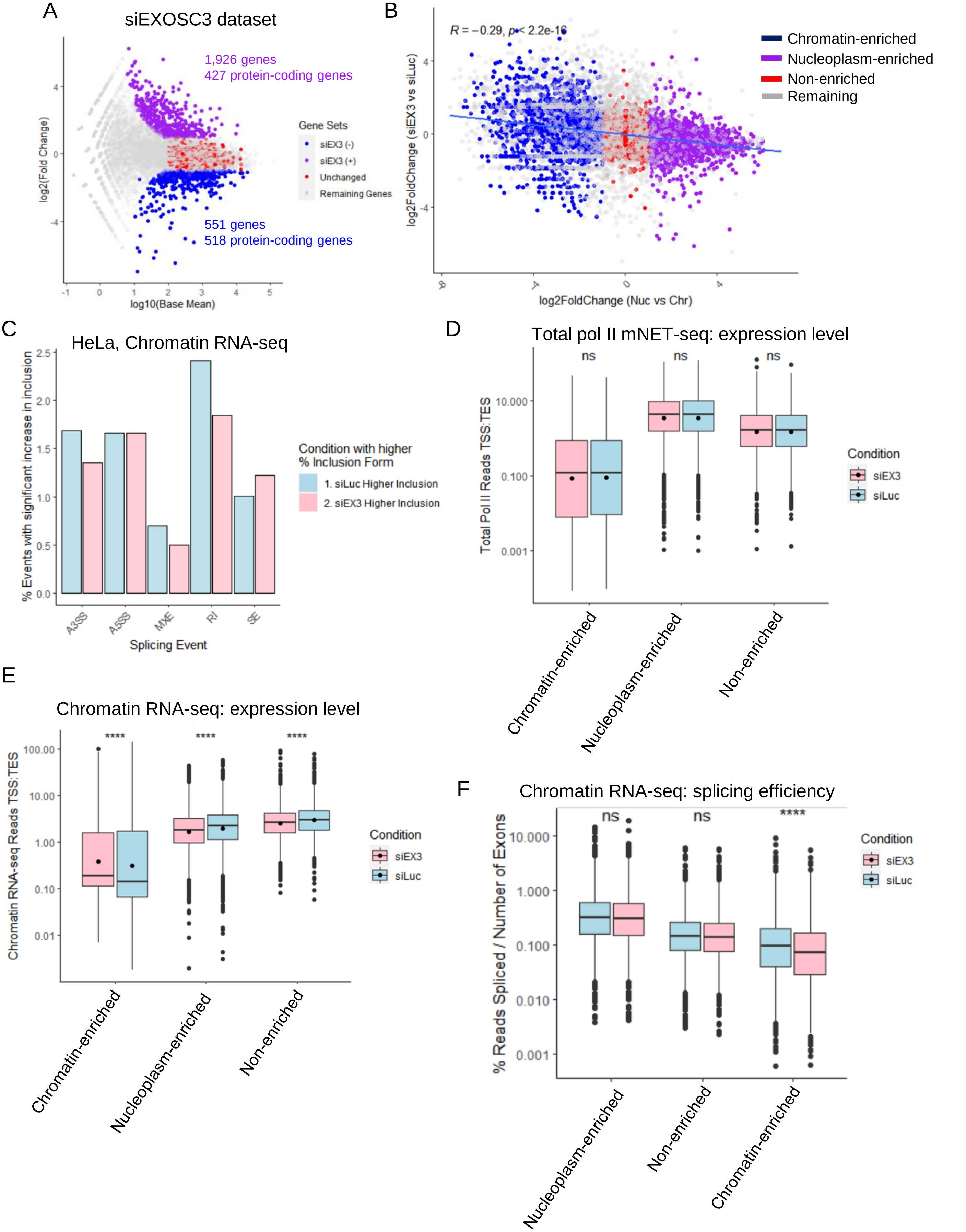


Supplementary Figure 7

**Supplementary Figure 7. Transcripts from chromatin-enriched gene are sensitive to the nuclear RNA exosome**.

**(A)** MA plot in HeLa cells of the intron-containing protein-coding genes found to be upregulated (siEX3(+), purple) or downregulated (siEX3(-), blue) after siEXOSC3. A set of non-enriched genes (red) and the remaining genes (grey) are also indicated. **(B)** XY correlation plot of the nucleoplasm fold enrichment, defined as the log2 fold change between nucleoplasm RNA-seq versus chromatin RNA-seq, and the log2 fold change in siEXOSC3 versus siLuc. Transcripts from chromatin-enriched, non-enriched, and nucleoplasm-enriched genes are shown in blue, red, and purple, respectively. The Pearson correlation with p-value is indicated on the plot. **(C)** Bar charts of significant changes in chromatin RNA-seq of splicing events obtained with rMATs in control (siLuc, blue) or following the knockdown of the nuclear RNA exosome (siEX3, pink). A3SS: alternative 3' splice site; A5SS: alternative 5' splice site, MXE: mutually exclusive exons; IR: intron retention; SE: skipped exon. **(D) and (E)** Boxplots, shown as min to max with first quartile, median, and third quartile, of the expression level of full-length transcripts, including exons and introns, from the total pol II mNET-seq **(D)** and chromatin RNA-seq **(E)** data in HeLa cells in control (siLuc, blue) or after siEXOSC3 knockdown (siEX3, pink) of all nucleoplasm-enriched, chromatin-enriched, or non-enriched genes. Statistical test: Wilcoxon rank sum test. P-value: n.s. not significant, **** < 0.0001. **(F)** Boxplots, shown as min to max with first quartile, median, and third quartile, of the splicing index of each transcript normalized to the number of exons from the chromatin RNA-seq data in HeLa cells in control (siLuc, blue) or after siEXOSC3 knockdown (siEX3, pink) of all nucleoplasm-enriched, chromatin-enriched, or non-enriched genes. Statistical test: Wilcoxon rank sum test. P-value: n.s. not significant, **** < 0.0001.
